# Single-cell atlas of major haematopoietic tissues sheds light on blood cell formation from embryonic endothelium

**DOI:** 10.1101/774547

**Authors:** Maya Shvartsman, Polina V. Pavlovich, Morgan Oatley, Kerstin Ganter, Rachel McKernan, Radvile Prialgauskaite, Artem Adamov, Konstantin Chukreev, Nicolas Descostes, Andreas Buness, Nina Cabezas-Wallscheid, Christophe Lancrin

## Abstract

The Yolk Sac (YS) and Aorta-Gonad-Mesonephros (AGM) are two major haematopoietic regions during embryonic development. Interestingly, AGM is the only one generating haematopoietic stem cells (HSCs). To identify the difference between AGM and YS, we compared them using single-cell RNA sequencing between 9.5 and 11.5 days of mouse embryonic development and identified cell populations using CONCLUS, a new computational tool. The AGM was the only one containing neurons and a specific mesenchymal population, while the YS major component was an epithelial population expressing liver marker genes. In addition, the YS contained a major endothelial population expressing Stab2, a hyaluronan receptor, also highly expressed by liver endothelium. We demonstrated that the YS haematopoietic potential was restricted to Stab2-negative cells and that ectopic expression of Stab2 could reduce blood cell formation from endothelium. Our results indicate that the AGM is a tissue more favourable to HSCs development than the YS because of its microenvironment and the nature of its endothelial cells.

## Introduction

Haematopoietic Stem and Progenitor Cells (HSPCs) emerge from rare haemogenic endothelial cells (HECs) during embryonic development through the process called endothelial-to-haematopoietic transition (EHT) ^1–6^. This transition is known to occur in both large arterial vessels, most notably the Aorta-Gonad-Mesonephros (AGM) and the Yolk Sac (YS), at mid-gestation. Despite both tissues undergoing EHT to give rise to the burgeoning haematopoietic system, distinct differences have been noted in their contribution to embryonic haematopoiesis.

The AGM region of the mouse embryo has been identified as a key source of haematopoietic stem cells (HSCs) between embryonic (E) days 9.5 and 11.5 ^7^. These HSCs emerging from the AGM go on to colonise the foetal liver at E12 ^8^, where they proliferate before seeding the foetal spleen and bone marrow (BM) between E15.5 and E17.5 ^9^. From here, they will maintain blood cell production throughout the life of the organism.

In contrast to the AGM, the YS is an extra-embryonic membrane surrounding the embryo giving rise to many haematopoietic progenitor cells (HPCs) which lack the self-renewal characteristics of HSCs ^7^. One key function of the YS is the production of primitive erythroid cells to support the needs of the growing embryo from E7.25 ^10–12^. Additionally, it contributes to life-long haematopoiesis through the production of multi-potent Erythroid-Myeloid Progenitors (EMPs) from E8.25 ^10–12^, which give rise to definitive cell types including tissue-resident macrophages that persist into adulthood ^13, 14^. Although several studies suggested that HSCs may be produced from the YS^15–18^, engraftment experiments in avian models and transplantation studies in mice indicated an intra-embryonic origin of HSCs ^19–22^

To date no explanation has been found to account for the difference in output of EHT that occurs in the YS versus AGM. It is not known whether there are intrinsic differences between the HEC of the two tissues or whether cell-extrinsic factors produced in the microenvironment drive HSC output in the AGM. While the AGM has been studied intensely both at the cellular and molecular levels, comparatively little is known about EHT in the YS due to challenges associated with imaging this tissue and its lack of HSC potential. By comparing the process of EHT between these distinct tissues it could be possible to better understand what is essential to HSC production in the embryo.

A lack of markers identifying HEC has hampered efforts to understand EHT, however, single cell transcriptome analysis has enabled us to make advances towards understanding this rare cell type. Recent work has characterised haemogenic endothelial cells (HECs) of the AGM at the molecular level ^23^ whereas, the identity of endothelial cells with haemogenic capacity in YS is still unresolved. As such it is interesting to compare the similarities and differences of the haemogenic endothelial cells themselves and the microenvironment of these embryonic haematopoietic tissues. The microenvironment of the AGM has been more intensely studied than that of the YS. AGM mesenchymal cells and neuronal cells surrounding the aorta have been shown to induce EHT via the production of molecules such as catecholamines ^24^, BMP4 ^25–27^ and Hedgehog ^27^. Much less is known about signalling pathways between endothelial and microenvironment cells in the YS.

Given the precise timing and rare populations involved in EHT, single-cell RNA sequencing (sc-RNA-seq) is an ideal approach to identify and characterise the populations involved in the formation of the haematopoietic system. Recent single-cell atlases covered a broad spectrum of mouse embryonic development between E6.5 and E13.5 ^28, 29^. However, none of them addressed the differences of EHT between YS and AGM.

Here, we used sc-RNA-seq to investigate one specific key aspect of development, EHT from YS and AGM at E9.5 and E11.5. Analysis of sc-RNA-seq datasets requires sensitive methods for characterising both large clusters of dominant cell types and small populations in a tissue. Computational methods have been developed to perform clustering on sc-RNA-seq data ^30^. However, these algorithms were not entirely appropriate for our data. Consequently, we designed a new sc-RNA-seq pipeline called CONCLUS (CONsensus CLUStering, https://github.com/lancrinlab/CONCLUS) combining DBSCAN clustering algorithm with a Consensus Clustering approach relying on the sequential use of Principal Component Analysis (PCA) and t-Distributed Stochastic Neighbour Embedding (t-SNE) methods with multiple parameters to reach a reliable clustering solution.

CONCLUS allowed us to successful identify and annotate the distinct cell populations in the AGM and YS. Using this method, we identified clear differences between the YS and AGM in the types of cells that make up the microenvironment. Furthermore, although we identified key groups of endothelial cells with clear similarities between the two tissues, we also found a distinct difference in the presence of a dominant Stab2+ population in the YS. Moreover, expression of Stab2 is negatively correlated with haematopoietic potential and could play a role in the differential haematopoietic output observed in the AGM and YS.

## Results

### Single-cell RNA sequencing analysis of AGM and YS tissues

In an effort to better understand the previously described differences between YS and AGM haematopoiesis, we performed single-cell RNA sequencing at E9.5 and E11.5 (before and after HSC emergence in the AGM). Since endothelial cells represent a small population in these tissues, we enriched for cells expressing the endothelial marker VE-Cadherin (VE*-*Cad, also called *Cdh5*) and sorted VE-Cad Negative cells constituting the microenvironment (Fig. 1a). We then isolated single cells using the SMARTer ICELL8 Single-Cell System ^31^. We captured 1,284 cells at E9.5 and 1,482 at E11.5 and sc-RNA-seq was performed. For each run of sequencing, YS and AGM populations from the same time point were combined (Fig. 1b) to minimize any potential batch effect. After sequencing, the cell barcodes were used to assign each cell to its corresponding original condition according to the sorting strategy: AGM VE-Cad+, YS VE-Cad+, AGM VE-Cad-, and YS VE-Cad-.

**Figure 1:**
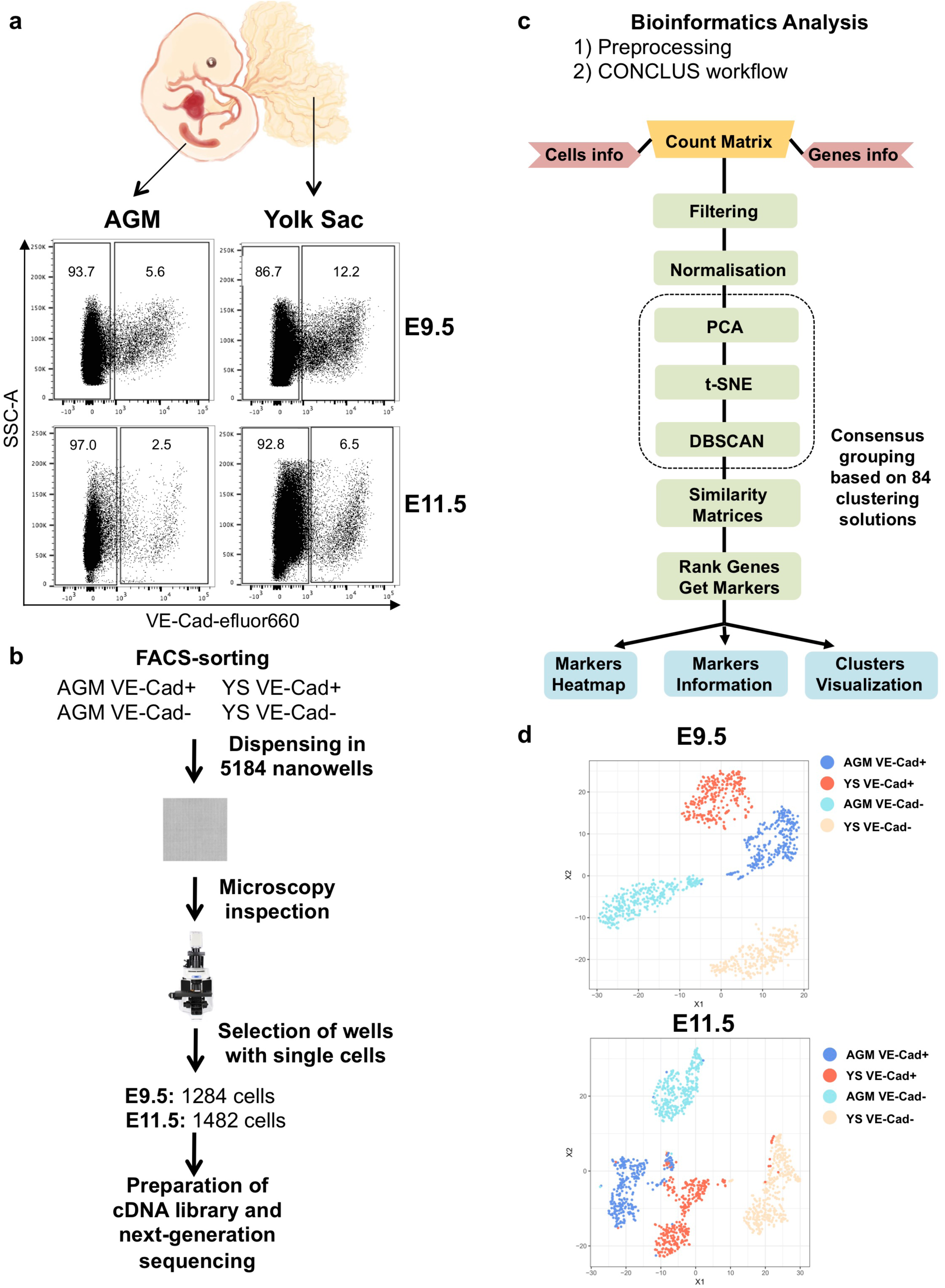
Workflow of sc-RNA-seq experiment to characterize endothelial and non-endothelial populations in haematopoietic sites of the mouse embryo. (**a**) FACS plot of cells isolated from AGM and YS regions at E9.5 and E11.5 showing VE-Cad expression versus SSC-A (internal complexity of the cells). (**b**) Experimental layout showing the process of sc-RNA-seq with the ICELL8 system. (**c**) Outline of the bioinformatics analysis. Dashed rectangle represents how CONCLUS generates 84 clustering iterations with DBSCAN. (**d**) t-SNE plots showing the cell distribution and their origin according to the initial sorting. Parameters: 8 PCs, perplexity 40.

We used several available bioinformatics tools ^32–35^ but they did not provide a robust clustering solution. For example, when using Seurat ^36^, a widely used sc-RNA-seq analysis tool, we noticed that the number of first principal components (PCs) and resolution could change the clustering. Therefore, numerous iterations were needed to identify stable clusters. The Hemberg lab proposed an approach overcoming this issue using an enumerative system of parameters enabling the definition of consensus clusters: SC3 ^32^. It simplifies the level of knowledge required for handling parameters. However, SC3 was not designed to explore rare populations, unlike GiniClust ^37^ and RaceID ^38^. Moreover, it is highly complex and not suitable for datasets with several thousand cells ^39^. Consequently, we reasoned that a tool taking advantage of the SC3 consensus idea, capable of processing thousands of cells and detecting rare populations could enable us to gain new insights into AGM and YS cellular composition.

We devised CONCLUS, an R package providing unsupervised clustering solutions to distinguish large clusters from rare populations accurately. By combining the consensus approach with the DBSCAN algorithm ^40^, we were able to identify sub-clusters differentiating AGM and YS environments. Following normalisation, we performed PCA and used seven ranges of PC and two values of perplexity to generate 14 t-SNE plots. In addition, we selected three values for Epsilon and two different MinPts values within DBSCAN. This enabled us to calculate 84 clustering solutions (14 t-SNE plots x 3 epsilon values x 2 MinPoints values), generate a similarity matrix and identify the most consistent clustering pattern (Supplementary Figure 1a & 1b). We found seven clusters at E9.5 and eleven at E11.5 (Fig. 2a). We could identify marker genes (Supplementary Figure 1c & 1d, Supplementary Files 1 & 2) and infer the nature of the identified subpopulations based on the literature and recent mouse single cell atlases ^28^. Even if the major clusters were also found with the Seurat algorithm, some of the smaller ones such as the YS_EMP at E9.5 and the EMP and Macrophage groups at E11.5 could not be identified (Supplementary Figure 2) while CONCLUS could clearly reveal specific marker genes for each of them (Supplementary Figure 1c & 1d).

**Figure 2:**
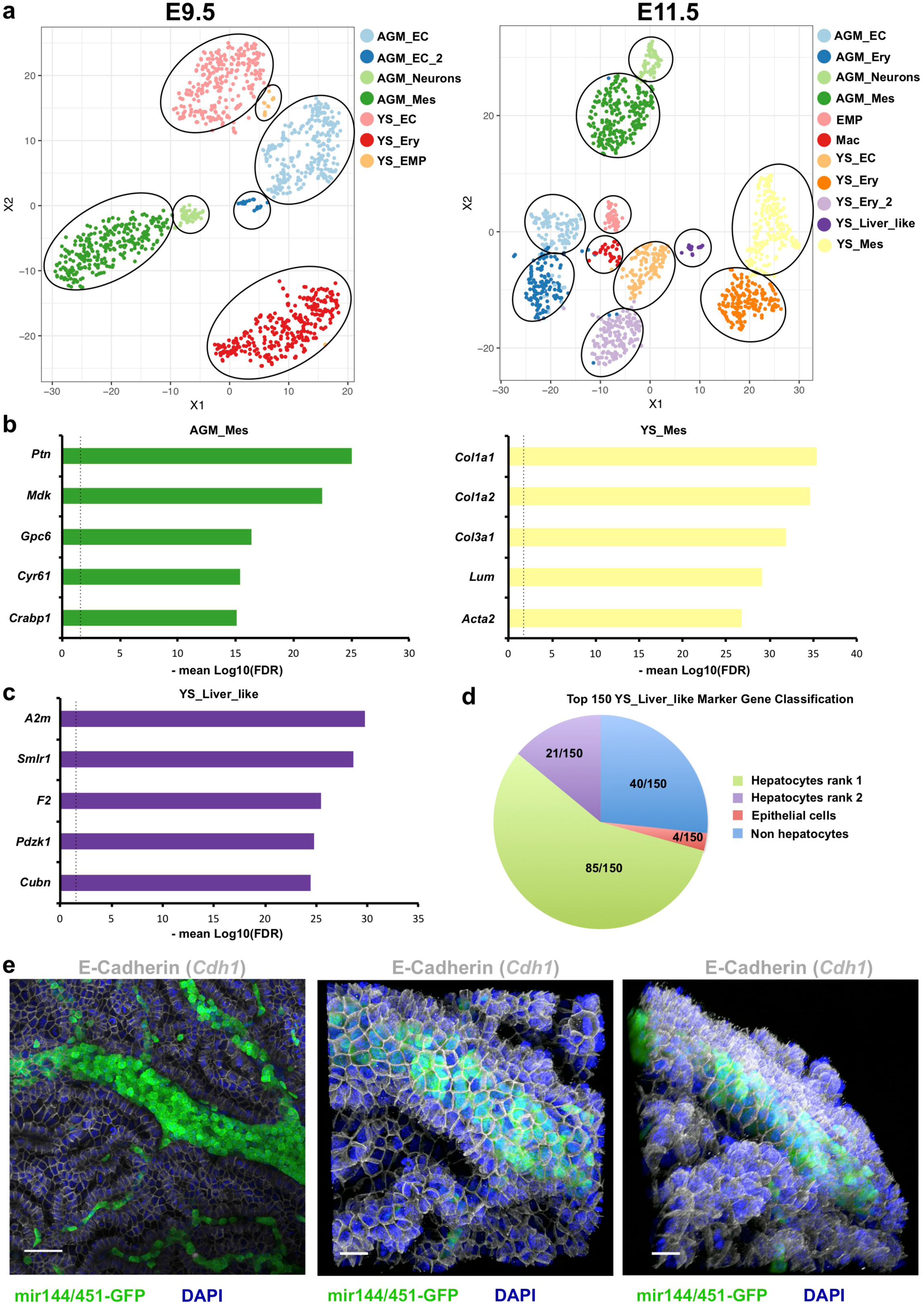
The YS contains a liver like population absent from the AGM. (**a**) t-SNE plots generated by CONCLUS showing the different cell clusters found at E9.5 (left) and E11.5 (right) by CONCLUS. (**b**) Bar plots showing the top 5 markers for the E11.5 AGM_Mes group (left) and the E11.5 YS_Mes cluster (right). The markers are ranked according to the -mean log10(FDR) of all paired comparisons between the target cluster and others. The vertical dotted line refers to the value 1.3 corresponding to a FDR of 0.05. (**c**) Bar plot showing the top 5 markers for the E11.5 YS_Liver_like group. The markers are ranked according to the -mean log10(FDR). The vertical dotted line refers to the value 1.3 corresponding to a FDR of 0.05. (**d**) Pie chart showing the classification of the top 150 YS_Liver_like Marker Genes according to the mouse organogenesis atlas. (**e**) Immunofluorescence analysis of E11.5 YS for the expression of miR144/451-GFP, E-Cadherin. The left panel corresponds to the 2D projection of the confocal microscopy. The middle and left show a 3D reconstruction of the confocal microscopy at different angles. Scale bars correspond to 50 μm.

### AGM and YS tissues have different microenvironments

The most striking difference between AGM and YS was at the level of non-endothelial cell populations. In AGM at E9.5 and E11.5, they were mostly mesenchymal (Mes) cells and neurons. In YS, at E9.5 non-endothelial cells were mostly primitive erythroid cells and at E11.5 they were a mix of primitive erythroid, mesenchymal and YS-liver like cells (Fig. 2a). The presence of mesenchymal cells and neuronal cells in the AGM has already been reported ^6, 24^ so we focused mostly on the clusters of YS microenvironment.

At E11.5 we found the mesenchymal populations of the AGM and YS (AGM_Mes and YS_Mes, respectively) to vary significantly with 1,154 differentially expressed genes (FDR < 0.05). At E11.5, AGM top 5 mesenchyme markers were *Ptn*, *Mdk Gpc6*, *Cyr61* and *Crabp1* while the YS top 5 were *Col1a1*, *Col1a2, Col3a1, Lum* and *Acta2* (Fig. 2b and Supplementary File 2). GO term analysis performed with DAVID Bioinformatics Functional Annotation tool ^41, 42^ showed that genes overexpressed by AGM_Mes were related to “nervous system development” (FDR= 2.36E-14), “axon guidance” (FDR= 4.75E-09) and “heart development” (FDR= 6.58E-06). In contrast, genes overexpressed by YS_Mes belonged to “collagen fibril organisation” (FDR= 8.41E-10), “endodermal cell differentiation” (FDR= 1.85E-04) and “angiogenesis” (FDR= 4.88E-04) (Supplementary File 1).

Only the AGM contained neuronal cells (AGM_Neurons). We detected them both at E9.5 and E11.5. Following GO term analysis of AGM Neuronal cell marker genes, we found the terms “nervous system development” (E9.5 FDR= 2.10E-15; E11.5 FDR= 5.30E-15), “axon guidance” (E9.5 FDR= 3.26E-12; E11.5 FDR= 4.11E-13) and “Wnt signalling pathway” (E9.5 FDR= 3.84E-06; E11.5 FDR= 7.66E-07) (Supplementary File 3).

At E11.5 we identified in the YS a small population whose top 5 marker genes were *A2m*, *Smlr1*, *F2, Pdzk1* and *Cubn* (Fig. 2c and Supplementary File 2). Interestingly, these five genes have been reported to be specific to the liver. To know whether these five genes were the only liver markers, we examined the top 150 marker genes of this population. We used the mouse organogenesis cell atlas covering development between E9.5 and E13.5 to investigate the specificity of these genes ^28^. This atlas offers the possibility to check the relative quantity of a given transcript across all the cell types detected in this developmental window. Out of these 150 genes, 85 (56.6%) were ranked as the most expressed in foetal hepatocytes, and 21 were listed in second place in terms of expression in hepatocytes in the mouse embryo (Fig. 2d). Top GO terms for this population were “cholesterol metabolic process” (FDR= 2.0E-13), “steroid metabolic process” (FDR= 4.5E-12) and “lipid metabolic process” (FDR= 1.85E-10) consistent with the key functions of the liver (Supplementary File 3). Overall, 70% of the top 150 genes of this YS population were highly expressed in foetal hepatocytes suggesting a high degree of transcriptional similarity of this YS population to hepatocytes. For this reason, we called it YS_Liver_like. We next decided to examine by microscopy how they were localised compared to the blood vessels. To visualise the YS_Liver_like population, we chose an antibody recognising one of its markers *Cdh1* coding for the protein E-Cadherin (E-Cadh) that is expressed by epithelial cells ^43^. We stained YS derived from miR144/451^+/GFP^ mouse embryos. MiR144 and miR451 are micro RNAs specific of red blood cells, both primitive and definitive ^44, 45^. GFP marked erythroid cells and allowed us to visualise easily blood vessels in the YS. To our surprise, E-Cadh-positive cells were very abundant in the YS and they surrounded the blood vessels from all sides (Fig. 2e) suggesting that they could have a significant influence on the vascular network.

### Similar myeloid populations were detected in AGM and YS

Among the identified clusters, we found five with haematopoietic features across the two time points. At E9.5, we detected erythroid cells only in the YS and then in both tissues at E11.5 (Fig. 2a, Supplementary Files 1 & 2). EMP appeared first in the YS at E9.5 while at E11.5 EMP and mature macrophages were present in both AGM and YS. Of note, these two myeloid cell groups were a composite of cells from the two tissues highlighting their similarity despite their different location (Fig. 2a, Supplementary Files 1 & 2). The macrophage subset was characterised by high expression of the *Mrc1* gene ^46^ coding for CD206. Other markers of importance were *Cx3cr1* ^47^, *Csf1r* ^48^ and *Sirpa* coding for SIRPα ^49^ the ligand of CD47, a potent inhibitor of phagocytosis ^50^. Interestingly, CD206+ macrophages expressing these genes were recently described to be important in the AGM to help the formation of HSCs ^51^.

### AGM and YS tissues have different endothelial cell populations

Blood cells come from the vasculature through the EHT process. Consequently, we examined next the endothelial cell populations residing in the AGM and YS tissues. Our study allowed us for the first time to compare endothelial cells from the AGM and YS at E9.5 and E11.5. Although they had genes in common such as *Cdh5*, *Kdr* and *Tek*, there were large differences in gene expression. At E9.5 there were three endothelial clusters: AGM_EC, AGM_EC_2 and YS_EC (Fig. 2a). Between YS_EC and AGM_EC, we found 1,213 differentially expressed genes (FDR < 0.05). Most of them (80.5 %) had a higher expression in the YS_EC group compared to AGM_EC and GO term analysis showed they belonged to the terms “GO:0030036∼actin cytoskeleton organization” (FDR= 2.39E-07), “GO:0001525∼angiogenesis” (FDR= 4.45E-07) and “GO:0006897∼endocytosis” (FDR= 4.31E-06) (Supplementary File 3).

The comparison of AGM_EC and YS_EC to the AGM_EC_2 group yielded similar lists of differentially expressed genes. The AGM_EC_2 expressed significantly more genes linked to the following GO terms: “GO:0006351∼transcription, DNA-templated”, “GO:0006355∼regulation of transcription, DNA-templated” and “GO:0045944∼positive regulation of transcription from RNA polymerase II promoter” suggesting that transcription is more active in these cells compared to the AGM_EC and YS_EC clusters (Supplementary File 3).

At E11.5 we identified only one EC cluster per tissue (Fig. 2a), and the number of differentially expressed genes went down to 380 (FDR < 0.05). Sixty-nine per cent of these genes were upregulated in YS_EC compared to AGM_EC and part of them belonged to the GO term “GO:0006898∼receptor-mediated endocytosis” (FDR= 1.13E-04) similarly to the comparison between YS_EC and AGM_EC at E9.5.

Interestingly, the YS endothelium consistently expressed more the *Stab2* and *Lyve1* genes at both developmental time points compared to the AGM endothelial cells (Fig. 3a, Supplementary Files 1 & 2). *Stab2* and *Lyve1* are two genes coding for receptors involved in hyaluronan endocytosis ^52, 53^. They are markers of foetal and adult liver endothelial cells ^54^ (Supplementary Fig. 2) and help the liver clear hyaluronic acid molecules from the extracellular space ^55^. Using flow cytometry, we showed that at E9.5, E10 and E11, Stab2 was expressed by the majority of endothelial cells in the YS while the AGM VE-Cad+ cells were almost all Stab2 negative (Fig. 3b). These experiments confirmed our sc-RNA-seq analysis. In addition, microscopy analysis of Stab2 protein expression demonstrated that it was indeed present on YS blood vessels (Fig. 3c).

**Figure 3:**
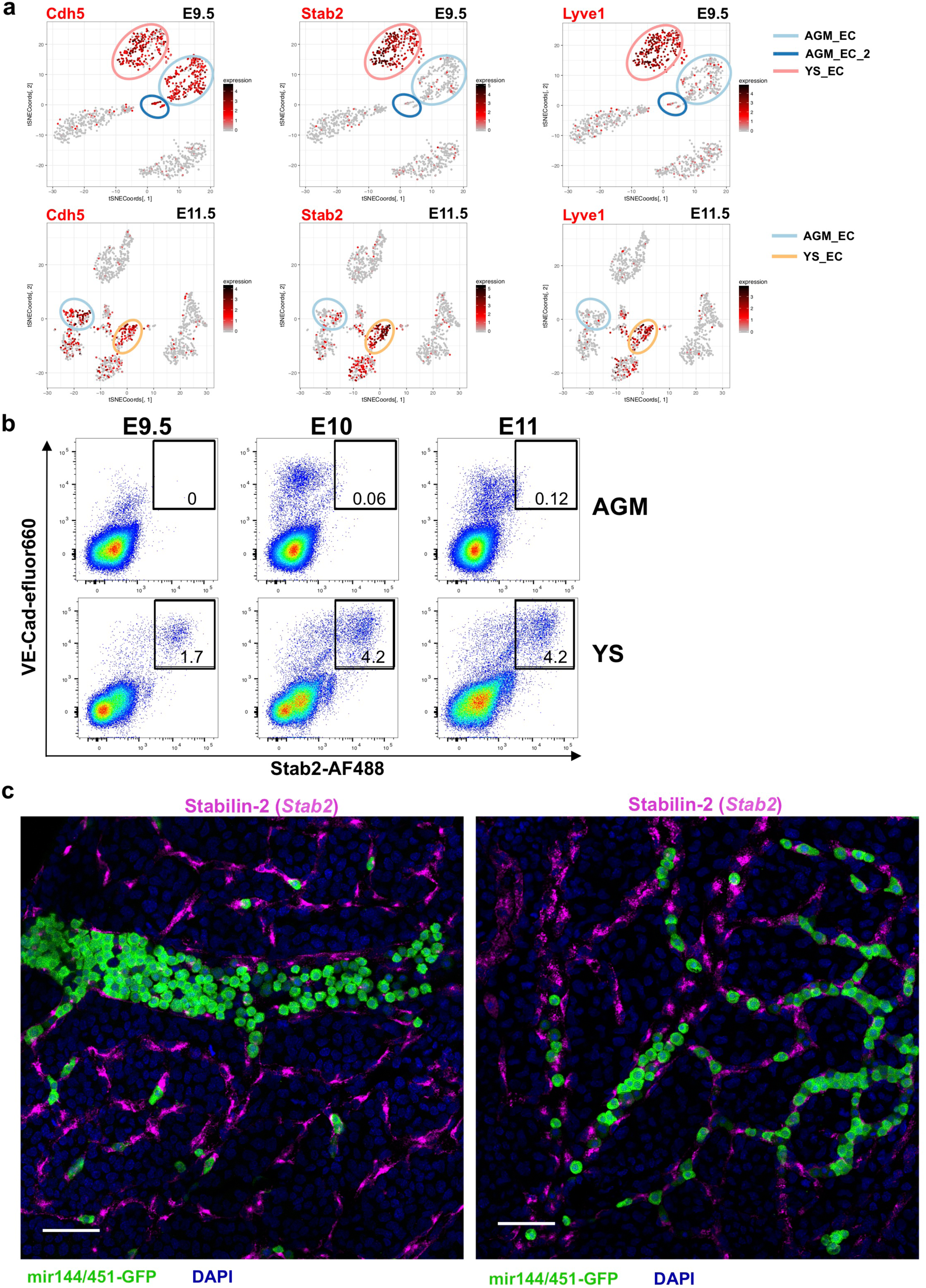
Yolk Sac and AGM have different types of endothelial cells. (**a**) t-SNE plots showing the expression of Cdh5, Stab2 and Lyve1 at E9.5 (top row) and E11.5 (bottom row). Cell coordinates are from Figure 2a. (**b**) FACS plot of cells isolated from AGM and YS regions at E9.5, E10 and E11 showing VE-Cad expression versus Stab2. (**c**) Immunofluorescence analysis of E11.5 YS for the expression of miR144/451-GFP and Stab2. Scale bars correspond to 50 μm.

### A fraction of YS endothelial cells co-expresses CD44 and Stab2

In a previous study, we demonstrated that in the AGM the haematopoietic capacity was restricted to VE-Cad^Pos^ cells expressing CD44, another hyaluronic acid receptor ^23^. The *Cd44* gene expression was low in our sc-RNA-seq dataset but it could be a limitation of the 3’ end sequencing technology that is less sensitive than full-length transcriptome analysis approaches. Consequently, we tested the expression of CD44 protein by flow cytometry (Fig. 4a). We found that some endothelial cells in the YS expressed this molecule. As in the AGM, we were able to distinguish three levels of expression among the VE-Cad+ compartment: CD44^Neg^, CD44^Low^ and CD44^High^. At E9.5, the majority of CD44^Neg^ and CD44^Low^ cells were Stab2^Pos^. At E10 and E11, only about half of CD44^Neg^ expressed Stab2 while the majority of CD44^Low^ cells remained Stab2^Pos^. Interestingly, the VE-Cad^Low^ CD44^High^ cells were not expressing Stab2 at any tested time points (Fig.4a). Using confocal microscopy, we confirmed the co-expression of Stab2 and CD44 proteins in the yolk sac vasculature (Fig. 4b - 4c). Following this finding, we decided to assess the haematopoietic properties of these CD44^Pos^ YS endothelial cells.

**Figure 4:**
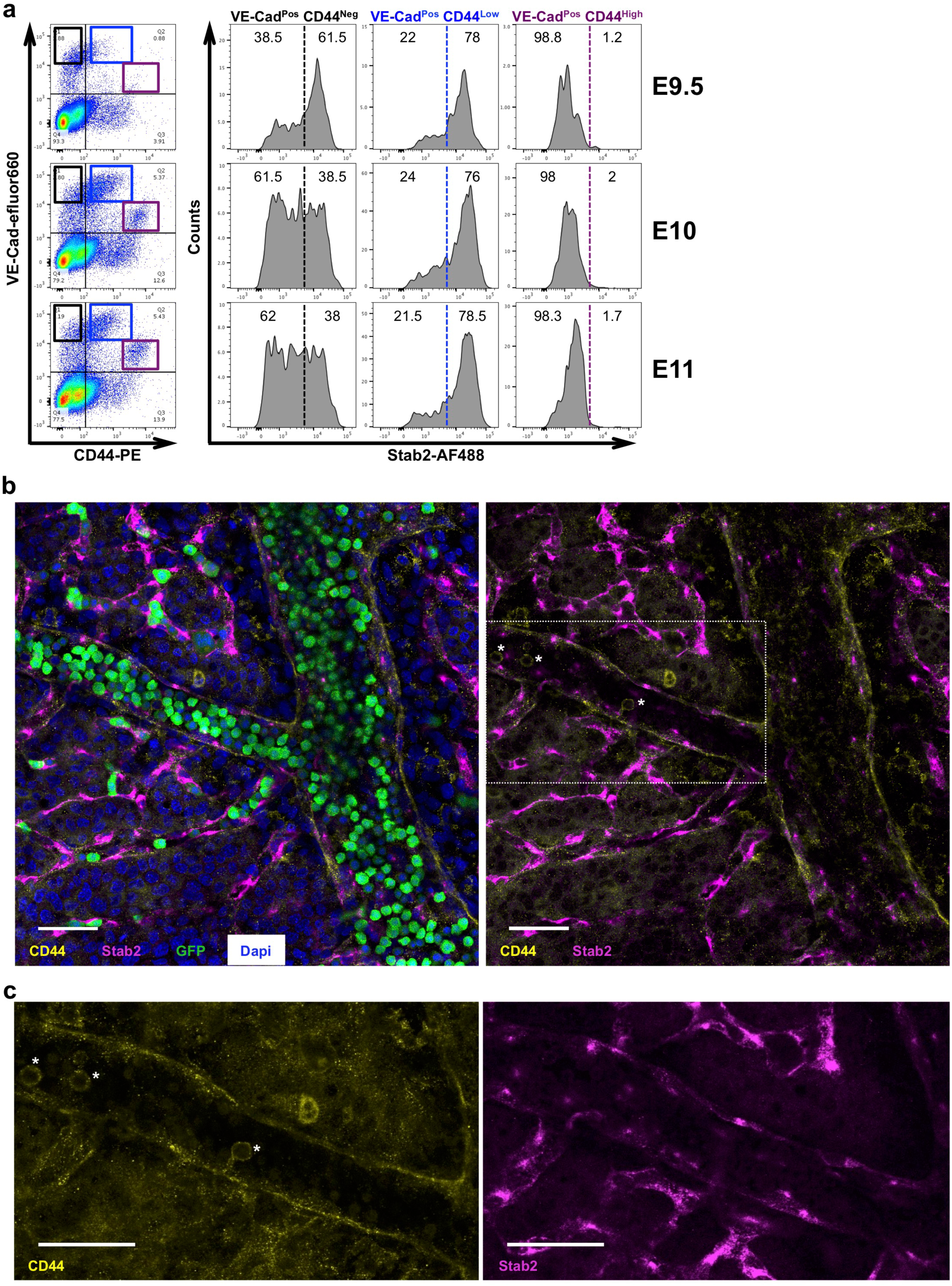
Hyaluronan receptors CD44 and Stab2 are co-expressed by endothelial cells in the YS. (**a**) FACS plot of cells isolated from YS at E9.5, E10 and E11 showing VE-Cad expression versus CD44 (left panel). On the right panel, histograms show the expression of Stab2 in the indicated populations. (**b**) Immunofluorescence analysis of E11.5 miR-144/451+/GFP YS for the expression of CD44, GFP and Stab2. White asterisks highlight blood CD44Pos cells in the lumen of the vessel. Scale bars correspond to 50 μm. (**c**) Enlargement of the area highlighted by the rectangle in (b). The left panel shows CD44 expression and the right shows Stab2. Scale bars correspond to 50 μm.

### Haematopoietic potential is enriched in the YS CD44^Pos^ Stab2^Neg^ endothelial fraction

In the AGM, we defined the haemogenic endothelial cells (HECs) as VE-Cad^Pos^ CD44^Low^ Kit^Neg^. These cells were characterised by the expression of *Grp126* (*Adgrg6*), *Pde3a*, *Sox6*, *Smad6* and *Smad7* ^23^. Pre-Hematopoietic Stem and Progenitor Cells (Pre-HSPC) type I were characterised by the VE-Cad^Pos^ CD44^Low^ Kit^Pos^ phenotype. They expressed endothelial and blood genes at the single-cell level. The Pre-HSPC type II population expressed Kit and VE-Cad at a lower level and CD44 at a high level. These cells expressed primarily blood genes and have down-regulated endothelial ones.

We asked whether or not we could find cells with HECs or Pre-HSPCs characteristics in the different fractions defined by the expression of Stab2 and CD44. At E10.5 and E11.5, we performed a single-cell FACS sort of the following populations: VE-Cad^Pos^ CD44^Neg^ Stab2^Neg^, VE-Cad^Pos^ CD44^Neg^ Stab2^Pos^, VE-Cad^Pos^ CD44^Low^ Stab2^Neg^ Kit^Neg^, VE-Cad^Pos^ CD44^Low^ Stab2^Pos^ Kit^Neg^, VE-Cad^Pos^ CD44^Low^ Stab2^Neg^ Kit^Pos^, VE-Cad^Pos^ CD44^Low^ Stab2^Pos^ Kit^Pos^ and VE-Cad^Pos^ CD44^High^. A total of 236 cells were subjected to single-cell q-RT-PCR using a panel of primers defined specifically to detect the HECs and Pre-HSPCs populations ^23^. Following clustering analysis, we found five clusters corresponding to two endothelial groups (YS_SC1 and YS_SC2), two co-expressing endothelial and blood genes (YS_SC3 and YS_SC4) and one expressing mostly haematopoietic genes (YS_SC5) (Fig. 5a).

**Figure 5:**
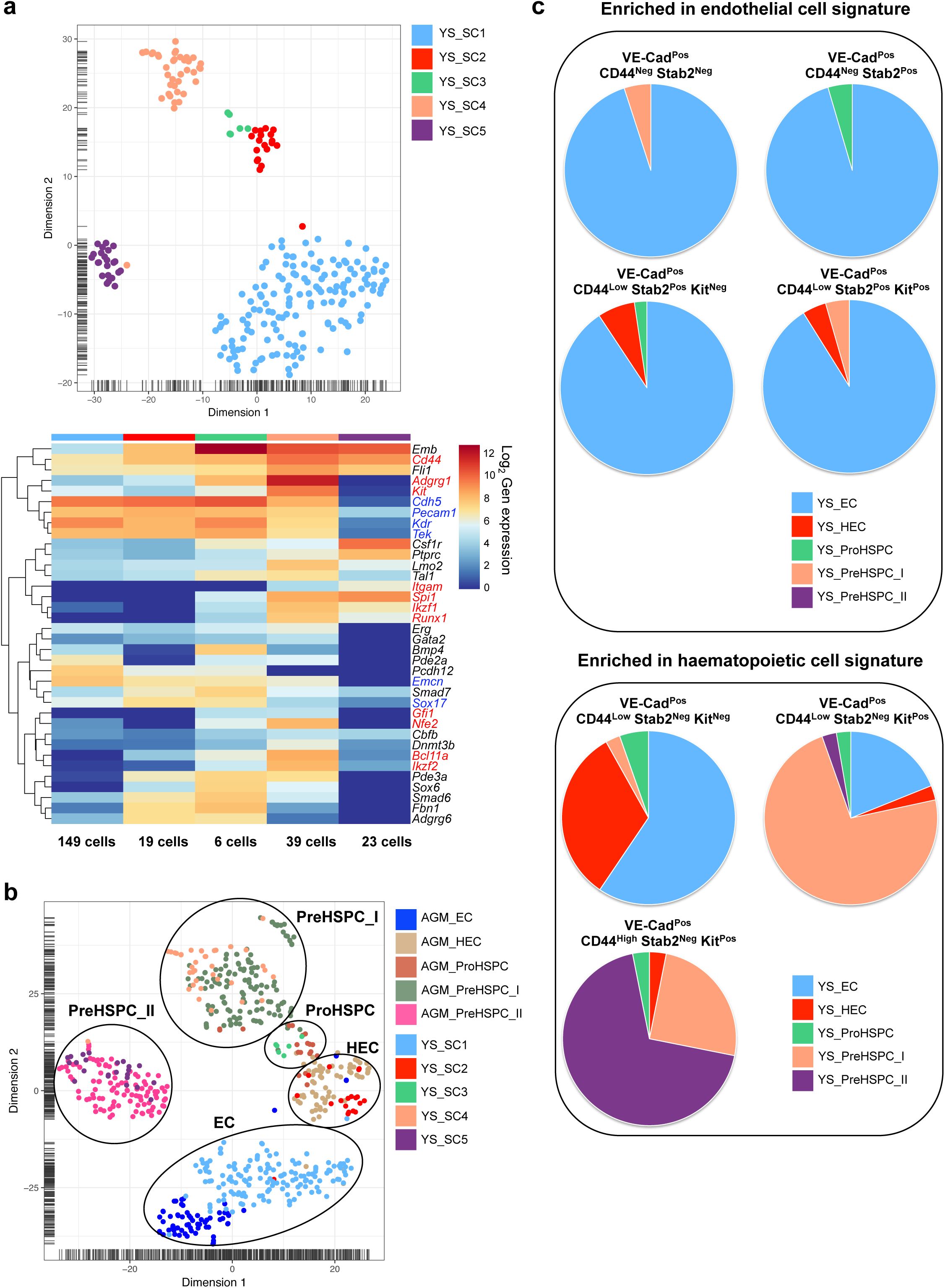
Haematopoietic transcriptional signature is enriched in the VE-Cad^Pos^ CD44^Low^ Stab2^Neg^ Kit^Pos^ and VE-Cad^Pos^ CD44^High^ Stab2^Neg^ fractions. (**a**) t-SNE plot showing the different clusters obtained from FACS isolated YS endothelial single-cells at E10 (2 litters) and E11.5 (1 litter) following sc-q-RT-PCR of 96 genes (top panel) and corresponding heatmap of gene expression (bottom panel). (**b**) t-SNE plot showing the different clusters after combining single-cells from AGM and YS. Ellipses highlight EC, HEC, ProHSPC, PreHSPC_I and PreHSPC_II groups. (**c**) Pie-charts indicating the contribution to the five different clusters defined in (a) for each indicated cellular phenotype.

To determine the nature of these populations, we compared them computationally to 386 VE-Cad^Pos^ cells from AGM ^23^. YS_SC1 cells were closer to the AGM_EC than any other populations while the YS_SC2 were clustering with AGM_HEC. The YS_SC3 group was mixing with the AGM_ProHSPC whereas the YS_SC4 cluster was clustering with AGM_PreHSPC_I. Finally, the YS_SC5 cells were the most similar to AGM_PreHSPC_II (Fig. 5b).

In summary, we found that most of the cells expressing Stab2 (Kit^Neg^ and Kit^Pos^) or lacking the expression of CD44 displayed a clear endothelial transcriptional signature (Fig. 5c). In contrast, cells with HEC, Pro-HSPC, Pre-HSPC_I and Pre-HSPC-II transcriptional characteristics were confined to the VE-Cad^Pos^ CD44^Pos^ Stab2^Neg^ populations (Fig. 5c).

This transcriptome analysis suggested that the Stab2^Neg^ VE-Cad^Pos^ CD44^Low^ Kit^Pos^ cells would more readily generate blood cells than the Stab2^Pos^ VE-Cad^Pos^ CD44^Low^ Kit^Pos^ cells. To test this hypothesis, we sorted 100 cells from each subset on OP9 stromal cells to perform a co-culture supporting the formation of blood cells. Three days following the initial culture on OP9, the wells containing haematopoietic growth were counted. Over seven independent experiments (43 wells in total), 90% of wells which received Stab2^Neg^ VE-Cad^Pos^ CD44^Low^ Kit^Pos^ cells gave rise to blood cells versus 8% of wells with Stab2^Pos^ VE-Cad^Pos^ CD44^Low^ Kit^Pos^ cells. These results supported the transcriptome analysis and showed that the highest haematopoietic potential was within the Stab2^Neg^ VE-Cad^Pos^ CD44^Low^ Kit^Pos^ population (Fig. 6).

**Figure 6:**
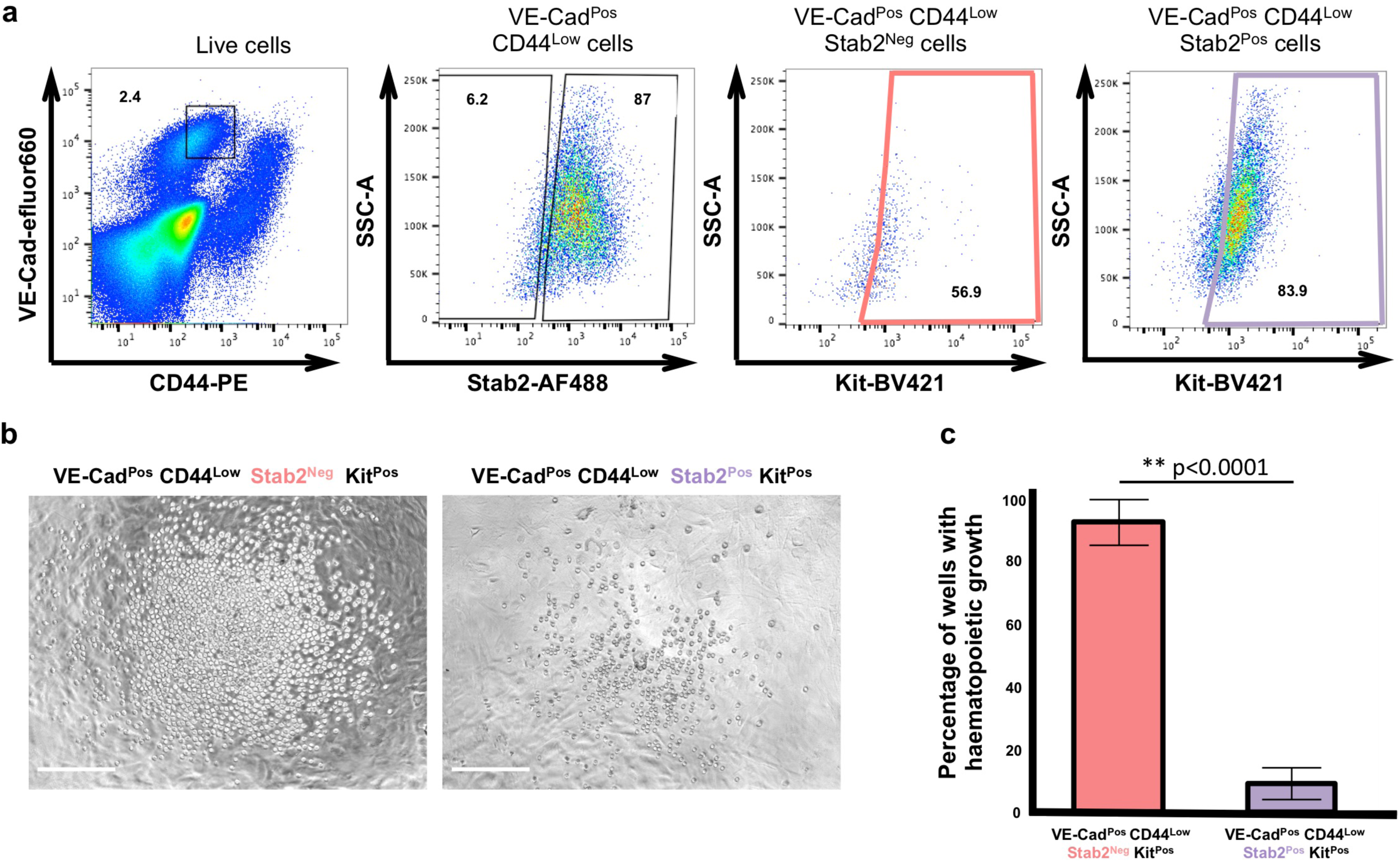
Haematopoietic potential is enriched in the VE-CadPos CD44Low Stab2Neg KitPos fraction. (**a**) Sorting strategy used to test the haematopoietic differentiation potential of VE-Cad^Pos^ CD44^Low^ Stab2^Neg^ Kit^Pos^ and VE-Cad^Pos^ CD44^Low^ Stab2^Pos^ Kit^Pos^ populations isolated at E10.5. (**b**) Images of haematopoeitic cell growth from 100 cells of VE-Cad^Pos^ CD44^Low^ Stab2^Neg^ Kit^Pos^ and VE-Cad^Pos^ CD44^Low^ Stab2^Pos^ Kit^Pos^ populations three days after OP9 co-culture. Scale bars represent 200 μm. (**c**) Bar graph showing the average percentage of wells (43 in total) in which haematopoietic cell growth has been detected for VE-Cad^Pos^ CD44^Low^ Stab2^Neg^ Kit^Pos^ and VE-Cad^Pos^ CD44^Low^ Stab2^Pos^ Kit^Pos^ populations. N=7, error bar represents standard deviation (p-value < 0.0001, two tailed t-test).

### Overexpression of Stab2 disrupts EHT *in vitro*

The fact that the most of the haematopoietic activity in the yolk sac was detected within the Stab2^Neg^ CD44^Pos^ compartment prompted us to ask whether or not *Stab2* expression could be a hindrance to the formation of blood cells. Previously, we have shown that disrupting the association between CD44 and Hyaluronan could interfere with the formation of blood cells ^23^. A total *Stab2* knock-out mouse model does not have any severe phenotype in relation to blood development suggesting that its presence is not essential for blood cell formation ^53, 56^. However, its expression on the surface of endothelial cells expressing CD44 might be a brake to blood cell generation.

To assess this possibility, we used the model of *in vitro* EHT based on the differentiation of embryonic stem cells (ESCs). In this model, a majority of endothelial cells produced by the haemangioblast express CD44 like in the AGM and also co-express *Smad6* and *Smad7* genes ^23^ but we did not know whether these cells were expressing Stab2 as well. Using flow cytometry, we tested the expression of this cell surface protein and we were surprised to find that ESC-derived ECs were Stab2^Neg^ (Supplementary Fig. 5a) unlike YS, where Stab2^Pos^ cells were the dominant endothelial population (Fig. 3). Considering that the ESC-derived ECs were Stab2 negative, they provided an excellent opportunity to test the effect of Stab2 overexpression on EHT.

To induce *Stab2* expression, we generated an inducible Stab2 (iStab2) ESC line where a transgene coding for a dead Cas9 fused to VP64-p65-RTA (dCAS9-VPR) transcriptional activators was under the control of the Tetracycline Response Element (TRE) promoter while a Stab2 specific guide RNA (gRNA) was expressed under a constitutively active promoter (Supplementary Fig. 5b). An ESC control cell line containing only dCAS9-VPR was also made. Following ESC differentiation into mesoderm, hemangioblast cells were sorted based on Flk1 expression and put in culture. We treated cells with doxycycline (dox) for three days and assessed the expression of VE-Cad and CD41 by flow cytometry. We verified that expression of dCAS9-VPR was induced following dox treatment in both lines (Supplementary Fig. 5c). As expected VE-Cad+ cells from the control line did not express Stab2 (Fig. 7a). In contrast, we could clearly detect a statistically significant increase of Stab2 expression with the iStab2 line even in the no dox condition suggesting that a low level of dCAS9-VPR expression was enough to trigger an upregulation of Stab2 expression (Fig. 7a). Addition of dox did not change significantly Stab2 expression. Of note, overexpression of Stab2 was mostly confined to VE-Cad+ cells (CD41+ and CD41-) (Fig. 7a).

**Figure 7:**
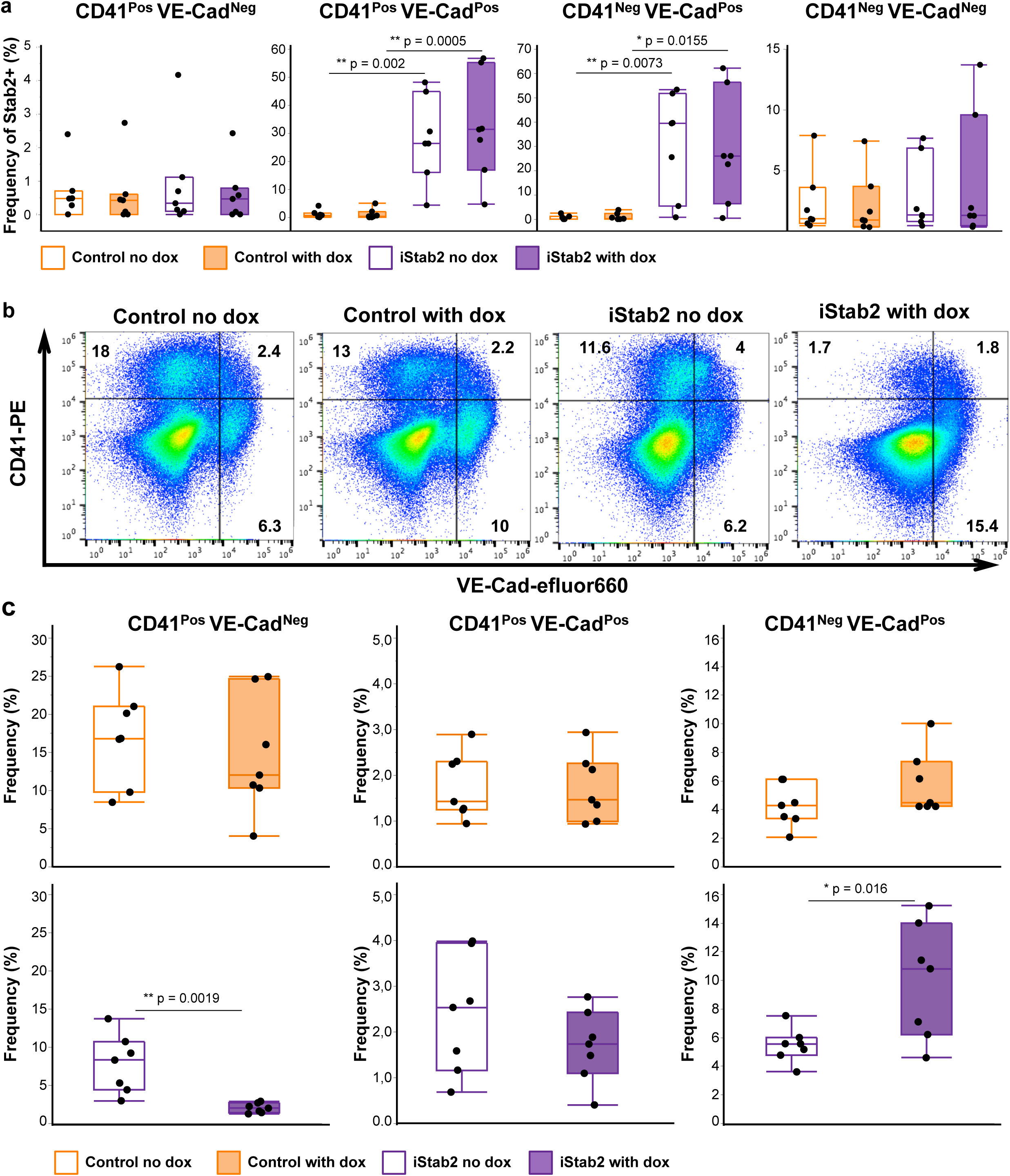
Overexpression of Stab2 disrupts in vitro EHT. (**a**) Tukey’s box plots showing the frequency of Stab2+ cells in the indicated populations. N=7 (* and ** indicate p-value < 0.05 and < 0.01 respectively, one-way ANOVA followed by Tukey’s multiple comparisons test). (**b**) FACS plot showing VE-Cad and CD41 expression after 72h of culture for the control and inducible Stab2 ESC lines in no dox and with dox conditions for one representative experiment. (**c**) Tukey’s box plots showing the frequency of CD41^Pos^ VE-Cad^Neg^, CD41^Pos^ VE-Cad^Pos^ and CD41^Neg^ VE-Cad^Pos^ populations for the indicated conditions. N=7 (* and ** indicate p-value < 0.05 and < 0.01 respectively, two tailed t-test).

When we compared the frequency of haematopoietic progenitors (CD41^Pos^ VE-Cad^Neg^) between the control and iStab2 lines in absence of dox, we noted a lower frequency of CD41^Pos^ VE-Cad^Neg^ cells in the iStab2 culture (Supplementary Fig. 5d). We obtained a similar result when we compared iStab2 no dox with iStab2 dox (Fig 7b - 7c). We also noticed an increase of the frequency of endothelial cells following dox treatment (Fig.7c). These results suggest that expression of Stab2 on endothelial cells could indeed slow down the EHT process.

## Discussion

In this study, we performed the first comparison of AGM and Yolk Sac cell composition between E9.5 and E11.5 of mouse embryonic development, the time window in which the definitive hematopoiesis is initiated. While both tissues are sites of embryonic haematopoiesis and undergo EHT, they contribute to the blood system in different ways. It remains unclear what are the similarities and differences between the endothelial cells of the vasculature and their surroundings that results in the differing output at these two sites. Using sc-RNA-seq, we compared the microenvironment and endothelium of each tissue to shed light on the process of EHT.

To analyse the resulting sc-RNA-seq datasets, we developed a consensus-clustering pipeline, CONCLUS, which allowed us to separate the different sub-populations in AGM and YS in more robust way compared to other well-known analysis methods.

Based on our sc-RNA-seq analysis, we found the YS and AGM to contain distinct populations within their microenvironments. The AGM is the only tissue containing neurons and a unique mesenchymal population, which have been involved in HSC generation ^24,25–27^. In contrast, the YS is the only tissue containing a cell population of extra-embryonic endodermal origin expressing many marker genes characteristics of foetal hepatocytes. In our sc-RNA-seq data, this population was very small but following immunofluorescence analysis, we found that it was a major component of the YS surrounding completely the vasculature. The discrepancy in population size between the transcriptome and microscopy analysis could lay in the preparation of the YS single-cell suspension for cell sorting. Indeed, epithelial cells tend to aggregate and they would be excluded through filtering and doublet exclusion performed during the FACS sorting procedure. This would explain the under-representation of this population in our sc-RNA-seq dataset. Overall, the abundance of this population revealed by confocal microscopy is likely to have a strong influence on the development of YS vasculature and the EHT process within it. It is supported by the phenotype of *Cubn* knock-out mouse embryos ^57^. *Cubn* is one of the top markers of this epithelial population and codes for a protein involved in the endocytosis of several ligands including vitamin B12 ^58^ and apolipoprotein A-I ^59, 60^. *Cubn* loss-of-function is embryonic lethal between E7.5 and E13.5. Mutant embryos have abnormal yolk sac where the epithelial cells derived from the visceral endoderm do not develop properly. As a result, the blood islands are abnormally large. In addition, there is a defect in the yolk sac vascular remodelling emphasizing the cross-talk between epithelial and endothelial cells ^57^. Indeed, after comparing the endothelium from AGM and YS, we identified significant differences in gene expression. Strikingly, the YS not only contains a liver-like epithelial population, but the majority of its vasculature expresses liver endothelial marker genes such as *Stab2*, *Lyve1* and *Mrc1* that are mostly absent from the AGM endothelium.

Following these results, we focused on the hyaluronan receptor Stab2, the most specific YS endothelial marker. We examined its expression in conjunction with CD44, another hyaluronan receptor. We have shown previously that there was a positive correlation between CD44 expression and haematopoietic identity of cells undergoing EHT in the AGM ^23^. Since we knew that co-expression of VE-Cad, CD44 and Kit was an excellent predictor of hematopoietic identity in the AGM, we aimed to find out if the same was true for YS. We found that a substantial fraction of endothelial cells in the YS expressed CD44. Unlike in the AGM endothelium, most of the CD44 positive cells of YS endothelium were also expressing Stab2. Of note, the expression of haematopoietic genes and the ability of producing cells were mostly restricted to the Stab2^Neg^ VE-Cad^Pos^ CD44^Pos^ Kit^Pos^ fraction. Therefore, the combined expression of VE-Cad, CD44 and Kit was not linked with blood potential in the YS if Stab2 was expressed. This result was consistent with an observation we made in a previous study ^69^. When we combined VE-Cad and the hematopoietic marker CD41 to isolate cells undergoing EHT in the YS, we found that only twenty-one percent of VE-Cad^Pos^ CD41^Pos^ cells were expressing blood genes. In contrast, sixty-five percent of VE-Cad^Pos^ CD41^Pos^ cells in AGM were expressing hematopoietic genes. This highlights the fact that markers commonly used in AGM to isolate cells undergoing EHT are insufficient to purify similar cells from YS unless Stab2 positive cells are excluded.

Our sc-RT-PCR results suggest similarities between AGM and YS regarding the populations involved in EHT. It is surprising considering the very different microenvironments that these tissues display. The expression of CD44 without Stab2 is specific of arterial endothelium, as we have shown previously ^23^. Since the YS also contains arterial endothelial cells belonging to the vitelline artery ^61^, this would explain our observations. It also suggests that the haematopoietic signature we observed in both AGM and YS CD44^Pos^ Stab2^Neg^ cells can arise independently of the microenvironment where the EHT takes place. However, the appropriate microenvironment is essential for the acquisition of HSC properties following EHT. Only the AGM has the proper cellular context for this process of maturation to occur.

Stab2 and CD44 are both hyaluronic acid (HA) receptors but they might antagonize each other if they are expressed together on the same cells due to their different modes of action. The role of HA in enhancing haematopoiesis was reported in multiple studies ^23, 62, 63^. We found that the interaction between CD44 and HA was beneficial to the EHT in the AGM and in the ESC based model of *in vitro* EHT. In contrast, Stab2 performs systemic clearance of HA from the extracellular matrix ^64^, which could reduce the interaction of CD44 with its ligand. The ectopic expression of Stab2 in our *in vitro* EHT model led to a significant reduction of haematopoietic output similarly to the disruption of CD44 binding to HA ^23^. Since the induction of Stab2 by dCAS9-VPR was restricted to VE-Cad+ cells, possibly because the Stab2 locus was more accessible in these cells compared to the other cell types in our culture, it suggests that the effect of Stab2 on EHT is affecting endothelial cells directly.

In conclusion, our work provides an important resource to understand the difference of haematopoietic outputs between AGM and YS. Moreover, it will pave the way to better define what is needed for endothelial cells to become blood cells. Finally, it further supports the importance of HA receptors at the onset of haematopoietic development. This knowledge will help further the development of cell culture methods to produce HSCs for therapeutic purposes.

## Materials and Methods

### Mouse lines and Embryo dissection generation

C57BL/6 and miR144/451^GFP/GFP^ two to three month-old mice were used for timed mating. The welfare of adult mice used in this work was covered by the licence n°17/2019-PR approved by the Italian Health Ministry. All experiments were performed following the guidelines and regulations defined by the European and Italian legislations (Directive 2010/63/EU and DLGS 26/2014, respectively). They apply to foetal forms of mammals as from the last third of their normal development (from day 14 of gestation in the mouse). They do not cover experiments done with day 12 mouse embryos and at earlier stages. Therefore, no experimental protocol or license was necessary for the performed experiments on mouse embryos. Mice were bred and maintained at the EMBL Rome Animal Facility in accordance with European and Italian legislations (EU Directive 634/2010 and DLGS 26/2014, respectively).

Embryos were obtained by timed mating between C57BL/6 wild-type mice or between miR144/451^GFP/GFP^ and C57BL/6 mice. Pregnant mice were killed by cervical dislocation between E9.5 and E11.5 of gestation. Uterine horns were collected; the maternal tissues were removed as well as the placenta to isolate the embryos. Embryos were collected in PBS supplemented with 10% FBS (PAA Laboratories). The yolk sac was torn gently and separated from the embryo proper by tearing off the umbilical and vitelline arteries. Then, the somite pairs of the embryos were counted to determine their developmental stage (E9.5-E11.5). To isolate the AGM, the head, tail, limb buds, ventral organs and somites were removed. Yolk sac and AGM samples from the same embryonic development stage were pooled together in the same tube. To generate single-cell suspension from the isolated tissues, AGM and yolk sac samples were digested with collagenase (Sigma C9722) for 30 minutes at 37°C, then stained as described below and taken to flow cytometry sorting.

### Flow cytometry sorting of AGM and YS for single-cell RNA sequencing experiments

Single cell suspensions of wild type AGM and YS at E9.5 and E11.5 were stained with rat anti-mouse CD144-efluor660 1:200 (eBioscience #50-1441-82, clone eBIOBV13) and 7AAD (Sigma, #A9400). Cells were analyzed and sorted on FACSAria (BD). After gating on cells and doublet exclusion, live 7AAD-cells were bulk sorted into 15ml falcons as VE-Cad^Pos^ endothelial cells and VE-Cad^Neg^ microenvironment cells, and taken for dispensing on ICELL8 chip (Takara), as described below.

We performed one experiment with E9.5 AGM/YS and one experiment with E11.5 AGM/YS. After flow cytometry sorting, VE-Cad^Pos^ and VE-Cad^Neg^ cells were stained with Hoechst 33342 (Cell Viability Imaging kit, Molecular Probes), and counted with the Moxi Z Mini Automated Cell Counter (ORFLO). Stained cell solution was diluted in a mix with diluent and RNase inhibitor (New England Biolabs) to 1 cell/50 nl for dispensing on the ICELL8-chip (Takara) with the MultiSample NanoDispenser (Takara). Positive (RNA from E14FF mouse embryonic stem cells) and negative controls were prepared according to the ICELL8 protocol and dispensed with the MSND into the respective nanowells of the chip. All nanowells of the ICELL8 chip were imaged with a fluorescence microscope (Olympus). The images were analysed with the CellSelect software (Takara). Live single cells positive for Hoechst-33342 were selected for lysis and reverse transcription inside the ICELL8 chip. RT reaction mix containing 5X RT buffer, dNTPs, RT e5-oligo (Takara), nuclease-free water, Maxima H Minus RT (Thermo Fisher Scientific) and Triton X-100 was prepared and dispensed into the previously selected nanowells with single cells inside. The chip was placed inside a modified SmartChip Cycler (Bio-Rad) for the RT reaction (42°C for 90 min, 85°C for 5 min, 4°C forever).

The cDNA of all single cells was collected together and further concentrated with the DNA Clean and Concentrator−5 kit (Zymo Research). The Exonuclease I (New England Biolabs) reaction of the cDNA (37°C for 30 min, 80°C for 20 min, 4°C forever) was performed inside a conventional thermal cycler. Afterwards, the cDNA was amplified with the Advantage 2 PCR Kit (Clontech Takara) containing buffer, dNTPs, Amp Primer (Takara), polymerase mix and nuclease-free water (95°C for 1 min, 18 cycles of 95°C for 15 s, 65°C for 30 s and 68°C for 6 min, followed by 72°C for 10 min and 4°C forever). The amplified cDNA was purified with Ampure XP Beads (Beckmann Coulter). The cDNA size distribution was obtained with the High Sensitivity DNA BioAnalyzer (Agilent) and quantification was performed with the Qubit (Life Technologies). Illumina library preparation was carried out by using Nextera XT DNA (Illumina). Tagmentation was performed in tagment DNA buffer, Amplicon Tagment Mix and 1 ng of purified cDNA (55°C for 5 min and 10°C forever), next Neutralize Tagment Buffer was added (room temperature for 5 min.). After incubation, the NexteraXT PCR reaction mix was prepared with Nextera PCR Mastermix, i7 Index Primer from the Nextera Index Kit (Illumina), Nextera Primer P5 and Tagmented cDNA-NT buffer mix (72°C for 3 min, 95°C for 30 s, 12 cycles of 95°C for 10 s, 55°C for 30 s and 72°C for 30 s, final 72°C for 5 min and 10°C forever). Ampure XP purification was performed with the finished library. The size distribution was checked on an Agilent BioAnalyzer. Samples were sequenced with the Illumina NextSeq 500 sequencer using the SMARTer ICELL8 Single-Cell System protocol (paired-end 3’ end sequencing with UMIs).

The first reads contained 11 nucleotides (nt) of a cell barcode for a cell and ten nt of UMI barcode for an RNA molecule. The second reads were the length of 130 nt and contained a piece of a gene from 3’ end. At E9.5, we sequenced 305 YS VE-Cad^Neg^, 326 YS VE-Cad^Pos^, 354 AGM VE-Cad^Neg^, and 299 AGM VE-Cad^Pos^ cells: 1284 cells in total. At E11.5, the numbers were 383 YS VE-Cad^Neg^, 388 YS VE-Cad^Pos^, 339 AGM VE-Cad^Neg^, and 372 AGM VE-Cad^Pos^ cells: 1482 cells in total.

### Single-cell RNA sequencing data pre-processing

Quality control of reads was done using the fastqc software. Then we separated reads by cell barcodes and saved them into one file per cell. Adapters were removed using cutadapt and alignment to the mouse genome mm10 (GRCm38.86) was done using STAR. We counted reads that overlapped exactly with one gene. Then for calculating Unique Molecular Identifiers (UMIs), we took into account the possibility of mutations in the UMI barcode caused by inaccurate sequencing. We assumed that two UMI barcodes with a difference of two mismatches could originate from one. If there was such a pair where one of UMI barcodes had only 10% of reads from another one, we joined those reads considering it as one UMI. In this way, we reduced the amount of false-positive UMIs. We created a UMI count matrix where columns were cells; rows were gene features. We did not include in a count matrix UMIs that had less than two reads. Later, we considered a gene to be expressed if it had at least one UMI.

### *S*ingle-cell RNA sequencing downstream analysis with CONCLUS

To perform cells quality check, we looked at the distribution across all cells of such features as the number of genes (i) per cell, the percentage of UMIs related to protein-coding genes (ii), the total number of UMIs (iii), and percentage of UMIs related to mitochondrial genes (iv). The first three metrics were considered ‘positive’, the last one – ‘negative’. For example, the more genes were detected in a cell, the better. On the opposite, an abnormally high percentage of mitochondrial counts in comparison to all other cells could mean that a cell was damaged and leaking. For each feature, we calculated the first (*Q1*) and the third (*Q3*) quartiles and the interquartile range (*IQR*). For the positive characteristics, we removed all cells with values below *Q1-1.5*IQR* and for the negative one above *Q3+1.5*IQR* that represent standard thresholds in boxplots for detecting outliers. This step allowed us to keep 1204 good quality cells at E9.5 (304 YS VE-Cad^Neg^, 288 YS VE-Cad^Pos^, 323 AGM VE-Cad^Neg^, and 289 AGM VE-Cad^Pos^) and 1311 cells at E11.5 (371 YS VE-Cad^Neg^, 336 YS VE-Cad^Pos^, 305 AGM VE-Cad^Neg^, and 299 AGM VE-Cad^Pos^). We stored values for filtering features together with a column for four sorting conditions in a metadata matrix.

After cells filtering, we selected only genes with more than ten total counts and created a SingleCellExperiment object containing a count matrix with UMIs and metadata. We computed factors with Scran ^65^ and used them for normalising the data ^66^. For calculating the factors, we used the following values for parameters: sizes = c(20, 40, 60, 80, 100) and clusters from a quickCluster function of Scran. Usually, if a cell has a factor equal or less than zero, it must be removed. But in our case, all factors were positive, so the number of cells did not change after normalisation. We deleted highly abundant haemoglobins having names starting with ‘Hba’ or ‘Hbb’ because they seemed to be the primary source of contamination in both datasets. Additionally, we excluded poorly annotated genes that did not have gene symbols to improve the clusters annotation process. It resulted in 11836 genes at E9.5 and 11434 at E11.5 that we used for clustering analysis and marker selection.

For calculating coordinates of two-dimensional t-SNE, we used the following ranges of principal components: 1:4, 1:6, 1:8, 1:10, 1:20, 1:40, 1:50, and two values of perplexity: 30 and 40 for both datasets. Then we created 84 clustering iterations with DBSCAN ^40^. For the E9.5 dataset, epsilon equal to 1.4, 1.6 and 1.8, and MinPoints 5 and 7 were applied. For the E11.5 data – epsilon 1.3, 1.4, and 1.5, MinPoints 3 and 4. We calculated a similarity matrix showing the fraction of clustering iterations were two cells were in one cluster. Consensus grouping was performed using a cutree function on the dendrogram of the similarity matrix. We explicitly asked the function to split the E9.5 dataset into eight groups and E11.5 set into 13. Two groups in E9.5 YS VE-Cad*+* were merged into YS_EC because the difference between them was explained by a slight gradient of gene expression. E9.5 YS_EC is indeed a group where cells have different transcriptional biases, however, not yet clear marker genes. At E11.5, we assigned a tiny, noisy group of cells to the closest cluster AGM_Ery, and additionally merged two almost equal sized groups into YS_Ery because they were similar to E9.5, that separation was supported only by the weak gradient of gene expression but not significant positive markers. Eventually, we obtained 7 clusters at E9.5 and 11 groups at E11.5 with distinct positive marker genes that were used for annotation.

To calculate the probability that a gene was a positive marker, we performed t-test with hypothesis “greater” for all possible pairs between a cluster of interest and all other groups. Note that the variance of a gene was estimated by taking all cells in the dataset, not only two compared clusters. For further ranking, we used FDR from these comparisons and information on how close two groups were to each other. We assumed that it is more probable to find a DE gene between two very distinct groups than between two similar groups. Thus we slightly raised the ranking of genes that allowed separating similar groups. The information about the distance between clusters we obtained from a similarity matrix of clusters. Similarity matrix of clusters is a simplified “bulk” version of the similarity matrix of cells where we calculated the median value for each group.

For a cluster k and a gene G, a score_G_ was defined in the following way:

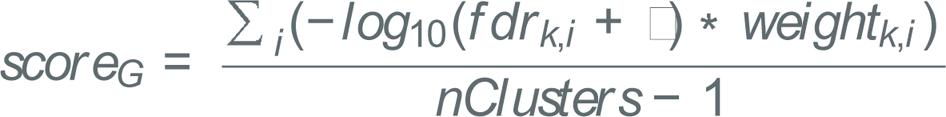

where

- *fdr_k,i_* is an adjusted p-value obtained by comparing expression of the gene *G* in cluster *k* versus expression of *G* in cluster *i*;
- *weight_k,i_* is a similarity between two groups. We defined it as the *<k,i>* element of the clusters similarity matrix plus *0.05* (to avoid zero scores in case if a group has zero similarity with others);
- *nClusters* is a number of consensus clusters;
- □ *=10^−300^* is a small number which does not influence the ranking and added to avoid an LJ error when *fdr* is equal to zero;
- k = [1,…,nClusters];
- *i = (*[*1,…,nClusters*] *except for* [*k*])

A gene with the highest score was assigned as the best positive marker for a cluster. To sum up, CONsensus CLUStering workflow (CONCLUS) (Fig. 1c, 1d, Sup. Fig. 3) includes such steps as cells and genes filtering, normalisation, similarity matrices of cells and clusters, consensus clustering, visualisation, and marker selection. The last step in CONCLUS is collecting information about marker genes from publicly available databases NCBI (citation), UniProt ^67^, and MGI ^68^ (Fig. 1c).

### Analysis of the Mouse organogenesis Atlas and Tabula Muris dataset

Seurat v.2 was used to perform sc-RNA-seq clustering analysis on the data retrieved from the Mouse Organogenesis Cell Atlas ^28^. We took the filtered data “cds_cleaned.RDS” from which the doublet and low-mRNA cells were removed. For annotation and metadata the “cell_annotation.csv” table was used.

We selected the cells related to the “Endothelial trajectory”. For these files we created Seurat object and ran clustering analysis including normalization with “LogNormalize” method. Afterwards, we found 4890 variable genes using the following parameters: x.low.cutoff = 0.0125, x.high.cutoff = 3, y.cutoff = 0.5. Next, we performed a scaling with linear model regressed on the number of UMI and clustering with dimensions 1:20 and resolution 1.0. With clustered data we generated t-SNE plots highlighting the genes we were interested in (Supplementary Figure 3).

The Tabula Muris Gene-count table for FACS sorted adult Liver cells was downloaded from Figshare: https://figshare.com/articles/Single-cell_RNA-seq_data_from_Smart-seq2_sequencing_of_FACS_sorted_cells_v2_/5829687.

The expression matrix was then processed through the CONCLUS pipeline with similar parameters to our AGM and YS datasets. We then generated t-SNE plots highlighting the expression pattern of *Cdh5*, *Stab2* and *Lyve1* genes (Supplementary Figure 3).

### Single-cell q-RT-PCR analysis

This experiment was performed exactly as described previously ^69^ using the Fluidigm Biomark system. Data analysis was also done as before ^69^.

### Yolk Sac flow cytometry sorting and clonogenic OP9 assays

Yolk sacs from E10-E10.5 embryos were dissected and dissociated as described above, then filtered through a 50μm sterile filcon (BD Biosciences #340630) to obtain a single cell suspension. The cells were counted and stained with the following rat anti-mouse antibodies: VE-Cad/CD144-efluor660 1:200 (eBioscience #50-1441-82, clone eBIOBV13); CD44-PE 1:2400 (BD Pharmingen #553134, clone IM7); CD117-BV421 1:200 (BD Horizon, #562609, clone 2B8); Stab2-AlexaFluor488 1:200 (MBL, #D317-A48, clone 34-2). Cells were analysed on FACSAria (BD) and sorted according to their VE-Cad, CD44, Stab2 and Kit expression (VE-Cad^Pos^ CD44^Low^ Stab2^Neg^ Kit^Pos^ and VE-Cad^Pos^ CD44^Low^ Stab2^Pos^ Kit^Pos^). Cell sorting was done onto OP9 cells growing in 96-well plates at 100 cells per well. Three days after sorting, wells were scored for “growth” or “no growth” of haematopoietic colonies. The percentage of wells with hematopoietic growth was calculated as the (number of wells with growth)*100/(number of total wells plated).

### OP9 growth and maintenance

OP9 cells were grown in alpha-MEM medium (Gibco #22561-021), containing 20% of fetal bovine serum (LGC, ATCC-30-2020). The day before yolk sac sorting, cells were replated into gelatinized 96-well plates at 3000 cells per well. On the day of sorting, existing OP9 medium was replaced with rich hemogenic endothelium mix, consisting of IMDM (Lonza BE12-726F); 10% fetal bovine serum (Gibco, #10270-42G9552K); 1% L-glutamine (Gibco, #25-030-024); human holotransferrin 0.6% (Roche, 10652202001); monothioglycerol 0.0039 % (Sigma, M6145); 25μg/ml L-ascorbic acid (Sigma, A4544); 0.0024 % of LIF 1 mg/ml (EMBL Heidelberg); 0.5% of recombinant murine SCF 10μg/ml (Peprotech, #250-03); 0.1% of recombinant murine IL-3 25 μg/ml (Peprotech, #213-13); 0.04% of recombinant murine IL-11 12.5μg/ml (Peprotech, #220-11); 0.1% of recombinant murine IL-6 10μg/ml (Peprotech #216-16); 0.1% of recombinant mouse Oncostatin M 10μg/ml (R&D Systems, #495-MO-025); 0.01% of recombinant human FGF basic protein 10μg/ml (R&D Systems, #233-FB-025).

### Immunofluorescence and confocal microscopy

The E11.5 Yolk sacs from miR144/451^+/GFP^ mouse embryos were first collected in 1xPBS+10% Fetal Bovine Serum (10270106, Gibco). Upon dissection they were fixed in 4% Paraformaldehyde solution in PBS (sc-281692, Santa Cruz) for 1hour at room temperature. After they were washed 3 times for 10min with 1x PBS, then incubated for 15min in 1xPBS with 20mM Glycine (G8898, SigmaAldrich). They were permeabilised for 30min in 1xPBS with 0.5% Triton-X100 (T8787, Sigma-Aldrich) at room temperature, blocked for 2 hours with a blocking buffer composed of 1xPBS, 0.1% Triton-X100, 2% BSA (A9418, SigmaAldrich) and 5% goat serum (S-1000, Vector Laboratories) at room temperature. Incubated with primary antibodies diluted in blocking buffer over night on a shaker at 4°C, rat anti-Stab2 1:100 (clone #34-2, MBL-D317-3) and rabbit anti-CD44 1:200 (polyclonal ab157107, Abcam).

The next day the Yolk sacs were washed 3 times for 10min with wash buffer composed of 1xTBS, 0.05% Tween20 (P9416, Sigma-Aldrich) and 2% BSA. The secondary antibody incubation was done over night at 4°C on a shaker, diluted in wash buffer, Alexa Fluor® 546 Goat Anti-Rat IgG (H+L) (A11081, Invitrogen) 1:800/1:400 and Alexa Fluor® 647 Goat Anti-Rabbit (A21244, Invitrogen 1:800/1:400).

Next day the Yolk sacs were washed 6 times for 15min with wash buffer and further 3 times for 10min washed with 1xPBS. Stained with DAPI (D1306, Invitrogen) diluted 1:1000 in 1xPBS incubated for 15min at room temperature, washed 3 times for 10min with 1xPBS. Finally they were mounted on Superfrost Plus slides (H867.1, Roth) and embedded in ProLong™ Diamond Antifade Mountant (P3697, Invitrogen) with a cover glass 22×40mm Nr1 (6310135, VWR). The slides were imaged on a Leica SP5 confocal microscope.

### Generation of ESC lines and in vitro ESC differentiation into blood and vascular lineages

A sgRNA (CCATCCCTAGTTGTTCCGCTAGG) was designed to target approximately 150bp away from the transcriptional start site of the mouse *Stab2* gene. The sgRNA was annealed with BstXI-BlpI overhangs for the insertion into a pPB_MU6_BFP plasmid (kind gift from Valentina Carlini, EMBL Rome). 80 μl of annealing buffer (10 mM Tris, pH 7.5-8.0, 60 mM NaCl, 1 mM EDTA) and 10 μl of 100 μM oligo1 (TTGCCATCCCTAGTTGTTCCGCTGTTTAAGAGC) and oligo 2 (TTAGCTCTTAAACAGCGGAACAACTAGGGATGGCAACAAG) each were used. Mixtures were heated at 95 °C for 3 min and then incubated at room temperature for 30 min.

A ligation reaction with sgRNA and linearised pPB_mU6_BFP was performed for 1h at room temperature using the reaction mix (annealed sgRNA 0.8 μl, vector 3.9 μl, buffer 2 μl, T4 ligase 1 μl, water 12.3 μl). Ligated sgRNA and plasmids were transformed into DH10B bacteria. After overnight incubation at 37°C, colonies were picked, miniprep was prepared using 5 NucleoSpin Plasmid/Plasmid (NoLid) protocol. Then a PCR reaction was performed using the forward primer (CTGCCCCGGTTAATTTGCAT) within the plasmid and the oligo 2 to detect if the ligation worked. For positive clones, an overnight midi-prep (50ml) with 1ml of culture was set up and DNA was purified using NucleoBond Xtra Midi/Maxi (Machery-Nigel, #740410.50).

To generate the iStab2 ESC line, A2loxEmpty ES cells ^69^ were transfected with a pPB_mU6_BFP plasmid (contains sgRNA and BFP), p118 PB-TRE-dCas9-VPR ^70^ containing dCas9 fused to VP64-p65-RTA (dCAS9-VPR) transcriptional activators and hygromycin resistance gene (Addgene, #63800) and a plasmid coding for the PiggyBac transposase (kind gift from Valentina Carlini). The transposase mediated the insertion of the two plasmids. To make the Control ESC line, only the p118 PB-TRE-dCas9-VPR and the transposase plasmids were transfected. The transfection was performed according to the manufacturer recommendations.

Twenty-four hours after the transfection, cells were treated with 200μg/ml HygromycinB (CalBiochem, #400051) in the morning and evening for two days and then only in the morning for 5 more days. After 7 days of antibiotic treatment, cells were FACS Aria sorted for BFP on 96-well plate of mouse embryonic fibroblasts. Fastest growing clones were selected and split until they were confluent for one well of a 6-well plate.

ES cells were then subjected to in vitro differentiation into Embryoid bodies to produce blood, endothelial and vascular smooth muscle cells following the same protocol described previously ^69^.

## Data availability

The raw count matrices and corresponding metadata for our sc-RNA-seq datasets can be downloaded here: https://oc.embl.de/index.php/s/NfKXoh6UI1Le1gW

## Supporting information

SupFile1

SupFile2

SupFile3

## Acknowledgements

We thank Cora Chadick (EMBL Rome FACS Facility, Italy), Giulia Bolasco (EMBL Rome Microscopy Facility, Italy), Paul Collier (EMBL Genomics Core Facility, Germany), Bianka Baying (EMBL Genomics Core Facility, Germany), Vladimir Benes (EMBL Genomics Core Facility, Germany), and Tallulah Andrews (Wellcome Trust Sanger Institute, UK) for advice and technical support. We are also grateful to Emmy Tsang (EMBL Rome), Georg Gasteiger (Institute of Systems Immunology, Würzburg, Germany) and Vsevolod Makeev, Sergey Bruskin, Artem Kasianov (members of the VIGG RAS Institute, Moscow, Russia) for helpful discussions.

## Conflict of interests statement

The authors declare no competing interests.

## Contributions

Maya Shvartsman, Polina V. Pavlovich and Morgan Oatley have made equally important contributions to this study and share first authorship. Maya Shvartsman, Conceptualization, Investigation, Formal analysis, Visualisation, Writing—review and editing; Polina V. Pavlovich, Conceptualization, Investigation, Software, Formal analysis, Visualisation, Writing—original draft, Writing—review and editing; Morgan Oatley, Conceptualization, Formal analysis, Investigation, Visualisation, Supervision, Writing—review and editing; Kerstin Ganter, Investigation, Formal analysis, Visualisation, Writing—review and editing; Rachel McKernan, Investigation, Visualisation, Writing—review and editing; Radvile Prialgauskaite, Investigation, Visualisation, Writing—review and editing; Artem Adamov, Formal analysis, Visualisation, Writing—review and editing; Konstantin Chukreev, Software, Formal analysis, Writing—review and editing; Nicolas Descostes, Software, Writing—review and editing; Andreas Buness, Supervision, Investigation, Writing— review and editing; Nina Cabezas-Wallscheid, Supervision, Investigation, Writing— review and editing; Christophe Lancrin, Conceptualization, Formal analysis, Supervision, Investigation, Visualisation, Methodology, Writing—original draft, Project administration, Writing—review and editing.

