## SupplementaryFigures for "Single-cell atlas of major haematopoietic tissues sheds light on blood cell formation from embryonic endothelium"

### Single-cell transcriptomics identifies the hyaluronan receptor Stabilin-2 as crucial for determining yolk sac haematopoietic output

Maya Shvartsman[1+], Polina V. Pavlovich[1,2,3+], Morgan Oatley[1+], Kerstin Ganter[1], Rachel McKernan[1], Radvile Prialgauskaite[1], Artem Adamov[1,3], Konstantin Chukreev[1,3], Nicolas Descostes[1], Andreas Buness[1,4], Nina Cabezas-Wallscheid[2] & Christophe Lancrin[1,*]

[1] European Molecular Biology Laboratory, EMBL Rome - Epigenetics and Neurobiology Unit, via E. Ramarini 32, 00015 Monterotondo, Italy.

[2] Max Planck Institute of Immunobiology and Epigenetics, Stübeweg 51, D-79108, Freiburg, Germany.

[3] Moscow Institute of Physics and Technology, Institutskii Per. 9, Moscow Region, Dolgoprudny 141700, Russia.

[4] Current address: Core Unit for Bioinformatics Data Analysis, Universitätsklinikum Bonn, Sigmund-Freud-Str. 25, 53127 Bonn, Germany

[+] Co-first authors

Supplementary Fig. 1

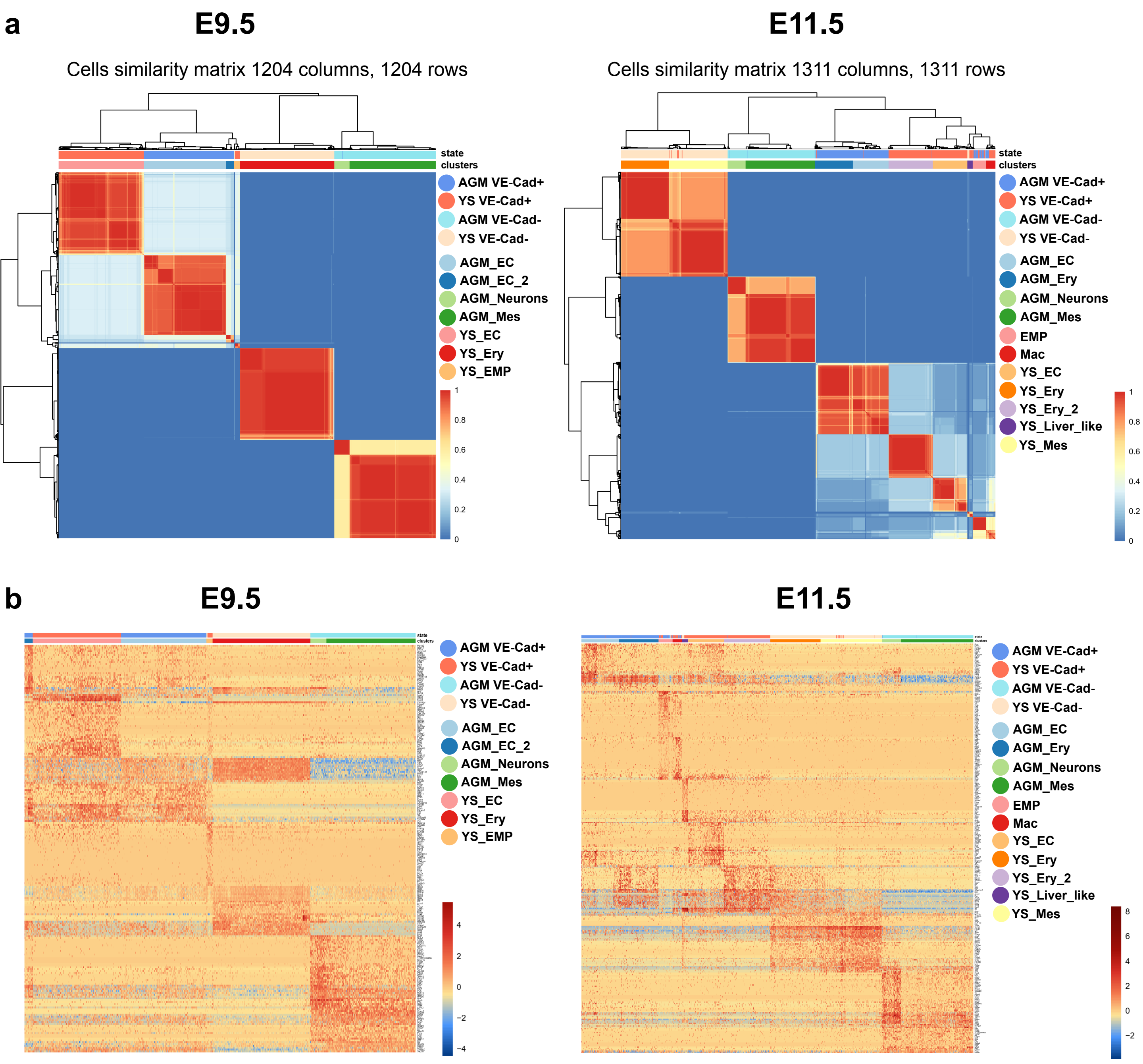


**Supplementary Fig. 1: CONCLUS cells similarity matrices and heatmaps with marker genes**

**(a)** Similarity matrices of cells for E9.5 (left) and E11.5 (right) datasets. **(b)** Heatmaps with top 35 positive marker genes per clusters. Mean-centred normalised data are shown. The legend for states and groups within an embryonic day is highlighted. Plots were generated with built-in functions of CONCLUS.

### Supplementary Fig. 2

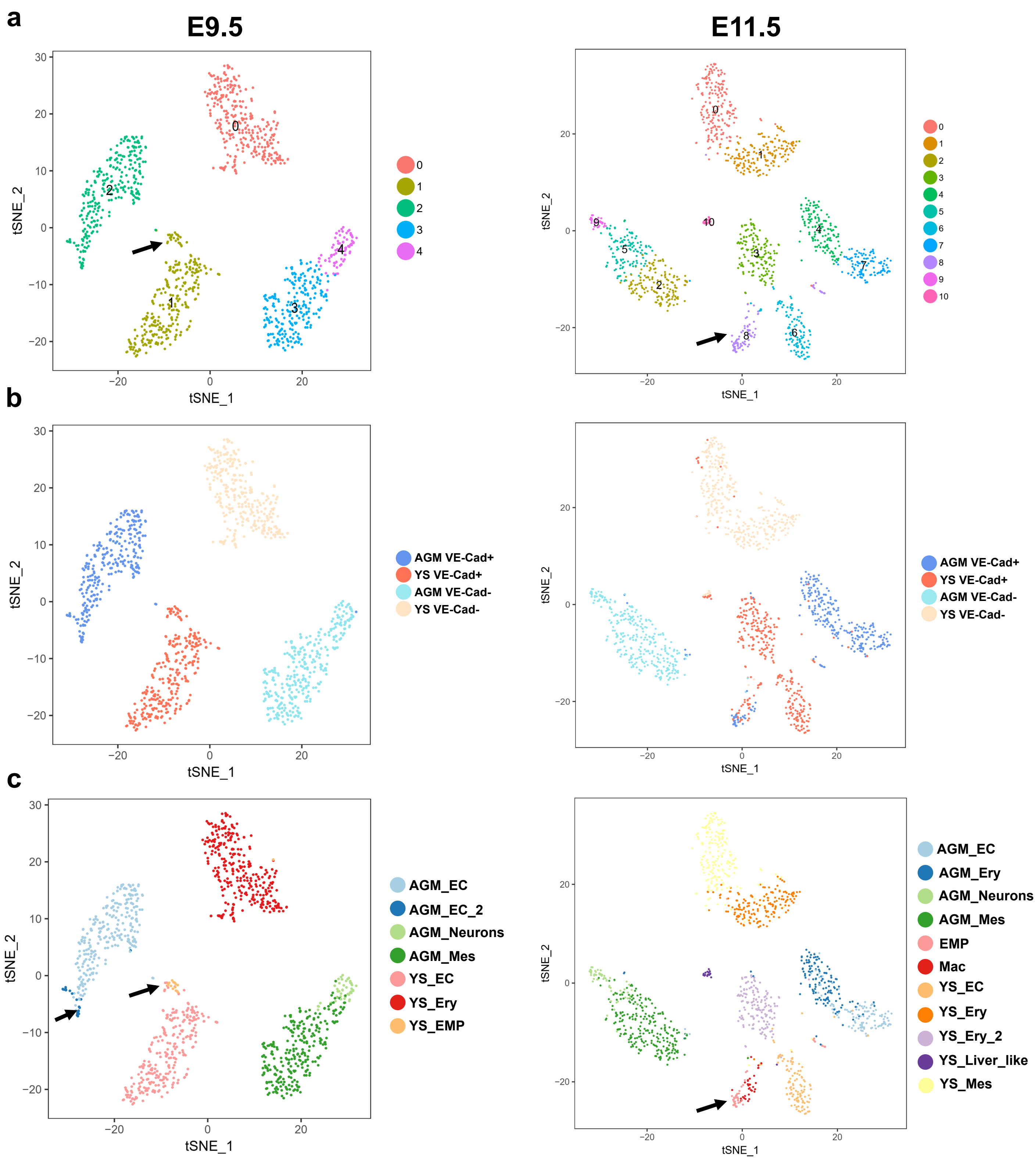


**Supplementary Fig. 2: Comparison of clustering of E9.5 and E11.5 datasets using Seurat and CONCLUS**

**(a)** t-SNE plots of E9.5 (left) and E11.5 (right) generated by Seurat and colored by clusters from Seurat. For E9.5, we chose scaling using negative binomial model and the resolution 0.6. Linear model regressed on the number of UMIs (nUMI) showed similar grouping. For E11.5, we show the result of the linear model regressed on the nUMI and the resolution 1.5. **(b)** t-SNE plots with the coordinates from (a) with highlighted states according to the design of the experiments. **(c)** t-SNE plots with the coordinates from (a) where colors show clusters of CONCLUS from the Figure 2a. The arrows indicate the clusters which were not found with Seurat but identified with CONCLUS.

### Supplementary Fig. 3

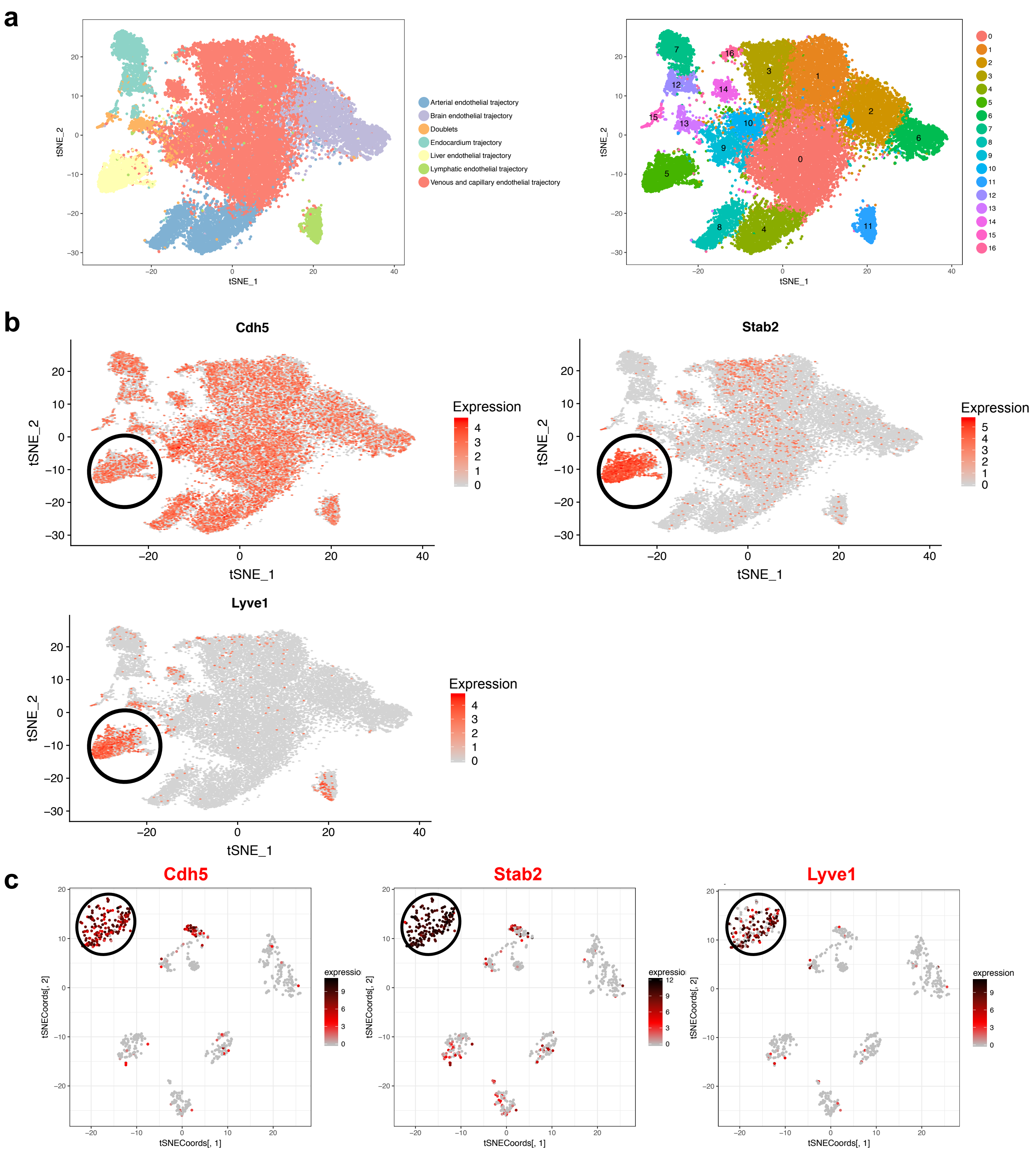


**Supplementary Fig. 3: Top YS endothelial marker genes are highly expressed by the liver endothelium in embryonic and adult stages**

**(a)** The t-SNE plots show the 26,107 endothelial cells from the Cao et al. mouse organogenesis atlas according to specific trajectories (left panel) and based on their belonging to distinct clusters (right panel). (**b**) The t-SNE plots show the expression of the indicated genes. The black ellipse highlights the cells with liver endothelial features: co-expression of *Stab2* and *Lyve1*. The scale of gene expression is indicated next to each plot. (**c**) The t-SNE plots show the liver single cells from the Tabula Muris dataset. The black ellipse highlights the cells with liver endothelial features: co-expression of *Stab2* and *Lyve1*. The scale of gene expression is indicated next to each plot.

#### Supplementary Fig. 4

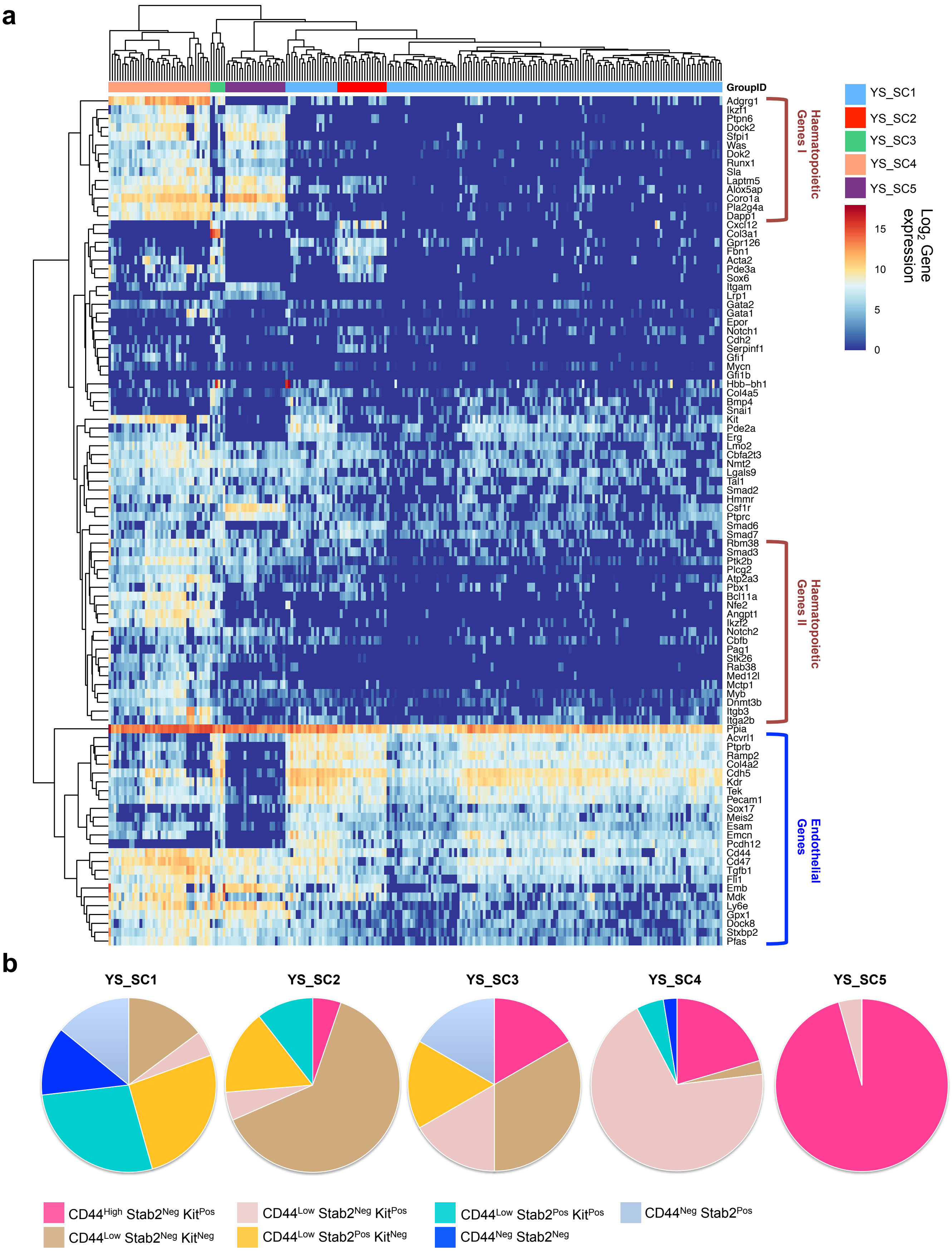


**Supplementary Fig. 4: Single-cell q-RT-PCR analysis of the different YS endothelial cell populations**

(**a**) Heatmap of gene expression showing the different clusters from FACS isolated YS single-cells of the indicated phenotypes at E10 and E11.5 following sc-q-RT-PCR of 96 genes. (**b**) Pie-charts indicating the phenotype of the cells composing the five different clusters defined in (a).

### Supplementary Fig. 5

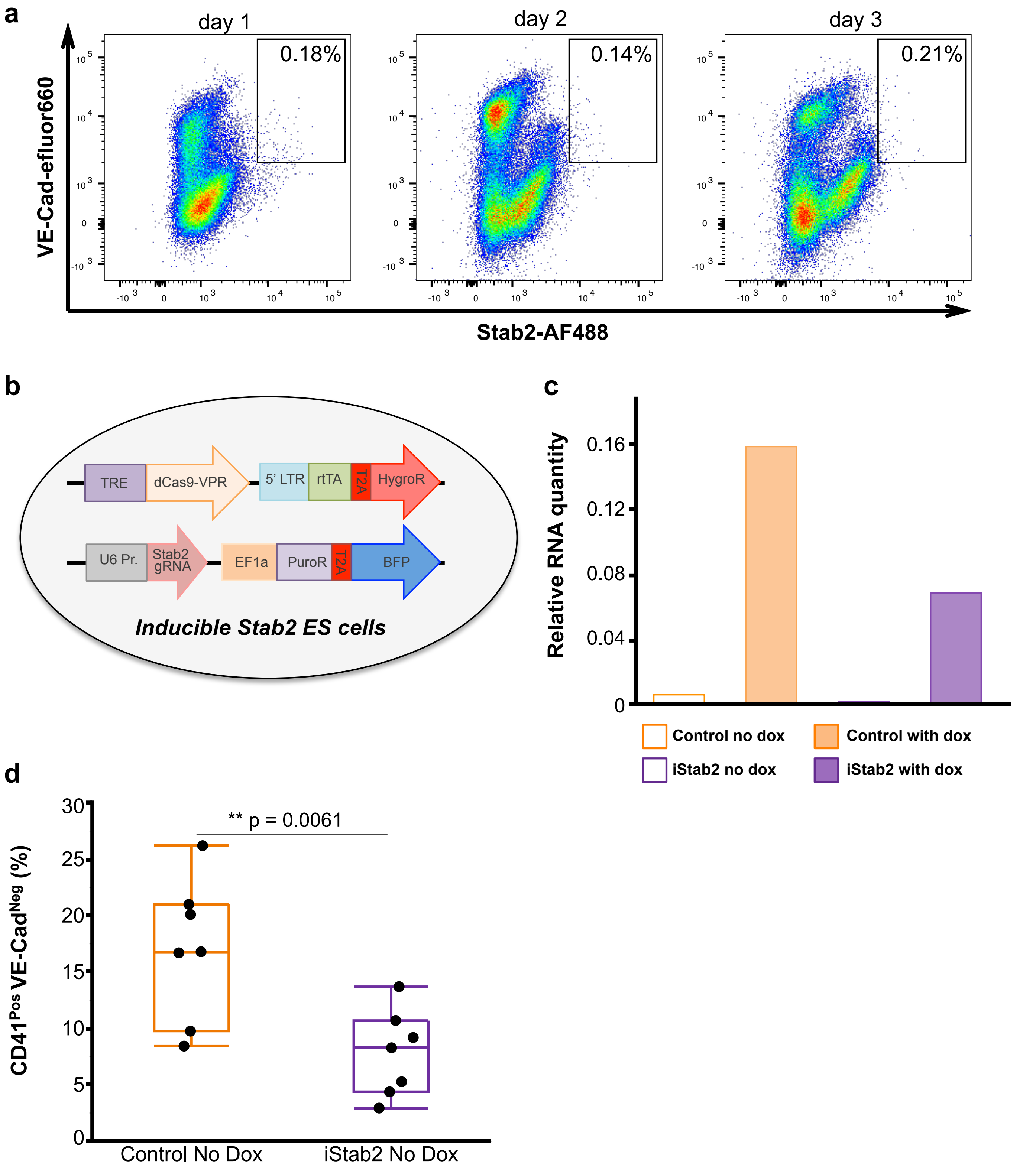


**Supplementary Fig. 5: Ectopic expression of Stab2 disrupts *in vitro* EHT**
(**a**) FACS plots showing the expression of VE-Cad and Stab2 in an *in vitro* haemangioblast differentiation time course. (**b**) Scheme describing the inducible Stab2 ESC line. (**c**) RT-q-PCR results for dCAS9-VPR at day 3 of haemangioblast culture for the indicated samples. A representative experiment is shown. (**d**) Tukey's box plots showing the frequency of $CD41^{Pos}$ $VE\text{-}Cad^{Neg}$ for the indicated conditions. N=7 (** indicate p-value < 0.01, two tailed t-test).

#### Description of the supplementary files

**Supplementary File 1: Marker Gene Lists of E9.5 clusters**
Each worksheet contains a list of top 100 marker genes for a given cluster. The genes are classified based on the statistical analysis performed by CONCLUS (see method section).

**Supplementary File 2: Marker Gene Lists of E11.5 clusters**
Each worksheet contains a list of top 100 marker genes for a given cluster (apart from the AGM_Ery cluster which had only 75 significantly up-regulated genes). The genes are classified based on the statistical analysis performed by CONCLUS (see method section).

**Supplementary File 3: Results of GO term analysis**
The worksheet contains the results of GO term analysis performed with DAVID Bioinformatics Functional Annotation tool on the marker gene lists generated by CONCLUS.
